# An oncogenotype-immunophenotype paradigm governing the myeloid landscape in genetically engineered mouse models of prostate cancer

**DOI:** 10.1101/2025.10.21.683668

**Authors:** Yuting Zhang, Yaoyin Li, Dailin Gan, Yan Liu, Xin Lu

**Author notes:** Correspondence: Xin Lu.

## Abstract

Distinct oncogenotypes sculpt divergent myeloid landscapes in prostate cancer. Using prostate-specific Ptenpc-/-, Ptenpc-/- Trp53pc-/-, Ptenpc-/- Smad4pc-/-, and Ptenpc-/- Trp53pc-/- Smad4pc-/- models, we identify a Smad4-loss-driven neutrophil-enriched subtype (NES) characterized by CXCL5, CXCL2, and CCL20 upregulation, polymorphonuclear myeloid-derived suppressor cell (PMN-MDSC) infiltration, and immune checkpoint resistance. In contrast, Smad4-intact tumors form macrophage-enriched subtypes (MES) responsive to immunotherapy. Mechanistically, Smad4 ablation activates YAP signaling and elevates histone epigenetic regulatory enzymes that enhance histone modification, chromatin accessibility, and transcription of neutrophil-recruiting cytokines. Genetic or pharmacologic inhibition of these enzymes suppresses chemokine expression, reduces neutrophil accumulation, restores CD8 T-cell activity, and limits tumor growth in immunocompetent hosts. Analyses of human prostate cancers support the findings from the murine NES prostate tumors. These findings establish a Smad4-YAP-epigenetic axis linking oncogenotype to immunophenotype and uncover a therapeutic vulnerability within the myeloid-dominant prostate tumor microenvironment.

## Introduction

The tumor microenvironment (TME) is a critical factor influencing both tumor progression and the efficacy of immunotherapies ^1–3^. Comprising a heterogeneous population of immune cells that varies among cancer types, the TME plays a complex role in either promoting or constraining tumor growth ^2^. Despite extensive studies on its immune and non-immune components, the precise regulatory mechanisms underlying their interactions and contributions to tumor dynamics remain inadequately understood. A deeper understanding of these processes is essential to develop more effective therapeutic strategies.

Among the immune cells within the TME, myeloid cells play a key role in modulating tumor immunity ^4–7^. Neutrophils and macrophages, known for their functional plasticity, utilize distinct mechanisms based on their activation state and local environment ^8–13^. Studies have classified tumors into neutrophil-enriched subtypes (NES) and macrophage-enriched subtypes (MES) ^14^. MES tumors, predominantly composed of CCR2-dependent macrophages, show variable responses to immune checkpoint blockade (ICB). In contrast, NES tumors display a systemic and local accumulation of immunosuppressive neutrophils or polymorphonuclear myeloid-derived suppressor cells (PMN-MDSCs), which are highly resistant to ICB. PMN-MDSCs are a subset of myeloid-derived suppressor cells (MDSCs), which represent a dominant immunosuppressive population in the TME across various cancer types ^15–17^. Further research is needed to investigate how tumor-intrinsic factors drive the preference for specific myeloid cell populations.

Prostate cancer (PCa) remains one of the most prevalent malignancies in men and a leading cause of cancer-related mortality worldwide ^18^. Its progression from localized disease to castration-resistant prostate cancer (CRPC) is driven by complex genetic and epigenetic changes, including mutations in key tumor suppressor genes such as *Pten* and *p53* ^19,20^. Notably, *Smad4* expression is significantly upregulated in *Pten^pc-/-^*prostatic intraepithelial neoplasia (PIN) compared to wild-type prostate epithelium, while downregulated *Smad4* expression has been observed in a subset of human primary prostate cancers and metastases ^21,22^. These alterations not only promote tumorigenesis but also reshape the TME, affecting the recruitment and function of immune cells. Recent studies have highlighted the role of tumor-infiltrating neutrophils (TINs)/PMN-MDSCs in driving immunosuppression and resistance to immunotherapy, thereby creating an environment conducive to tumor growth and metastasis in PCa ^23,24^. Targeting PMN-MDSCs has demonstrated potential in improving responses to immune checkpoint blockade (ICB) therapy in both primary and metastatic CRPC models ^24^.

To effectively target PMN-MDSCs, a comprehensive understanding of tumor-intrinsic molecular mechanisms is essential. Dysregulation of post-translational modifications (PTMs), such as methylation, acetylation, and phosphorylation, has been widely implicated in cancer progression ^25–27^. Among these PTMs, histone citrullination, mediated by enzymes of the Peptidylarginine Deiminase (PADI) family, is a key PTM regulating gene expression in cancer cells ^28,29^. Of particular interest are PADI2 and PADI4, both of which possess nuclear localization signals and play critical roles in modulating transcription. PADI4 has been shown to citrullinate histone H3R8, preventing the binding of HP1α and promoting the expression of pro-inflammatory cytokines such as IL-8 and TNF-α in cancer cells ^30^. Furthermore, PADI4 induces the citrullination of NF-κB, modulating the expression of IL-1 and TNF-α in neutrophils ^31^. Similarly, overexpression of PADI2 has been associated with increased tumorigenicity, epithelial-to-mesenchymal transition (EMT), and elevated levels of inflammatory cytokines (CXCL5, CXCL2, CXCL1, and IL-6) in breast cancer ^32,33^. Additionally, PADI2 has been shown to citrullinate MEK1, leading to the activation of downstream signaling pathways involved in tumor malignancy ^34^. Moreover, PADI2-mediated citrullination of histone H3R26 enhances the expression of androgen receptor (AR)-targeted genes in prostate cancer ^35^.

This study aims to investigate the role of PADI2 and PADI4, in regulating neutrophil recruitment and function within the prostate cancer TME. We demonstrate that genetic alterations, particularly the loss of *Smad4*, lead to the upregulation of PADI2 and PADI4, which promotes the recruitment of PMN-MDSCs and contributes to tumor progression. Through a combination of *in-vitro* and *in-vivo* approaches, we seek to elucidate the molecular mechanisms by which PADI2/4 influence the TME and evaluate the therapeutic potential of targeting these enzymes to enhance anti-tumor immunity.

## Results

### A Distinct Myeloid Cell Distribution Across Different Genotypes of Tumor Models

To elucidate the detailed immune cell composition of the tumor microenvironment (TME), we utilized flow cytometry to study key immune cell populations in prostate-specific spontaneous mouse models of prostate cancer (Probasin(PB)-Cre^+^lox mice). Major immune cell populations were analyzed when tumors reached a comparable size across four genotypes: *Pten^pc-/-^*(P), *Pten^pc-/-^ p53^pc-/-^* (PP), *Pten^pc-/-^ Smad4^pc-/-^* (PS), and *Pten^pc-/-^ p53^pc-/-^ Smad4^pc-/-^* (PPS) (Figure 1A). To collect tumor tissue, mice of each genotype were monitored until their tumors reached similar sizes (0.5–1 g) at varying ages (Figure S1A). Tumors in PS and PPS mice typically reached this size around 16 weeks, while tumors in PP and P mice required approximately 25 weeks and 45 weeks, respectively. In the survival comparison, mice bearing PS and PPS tumors exhibited significantly shorter survival times compared to those bearing PP and P tumors, indicating that PS and PPS tumors are more aggressive (Figure 1B).

**Figure 1:**
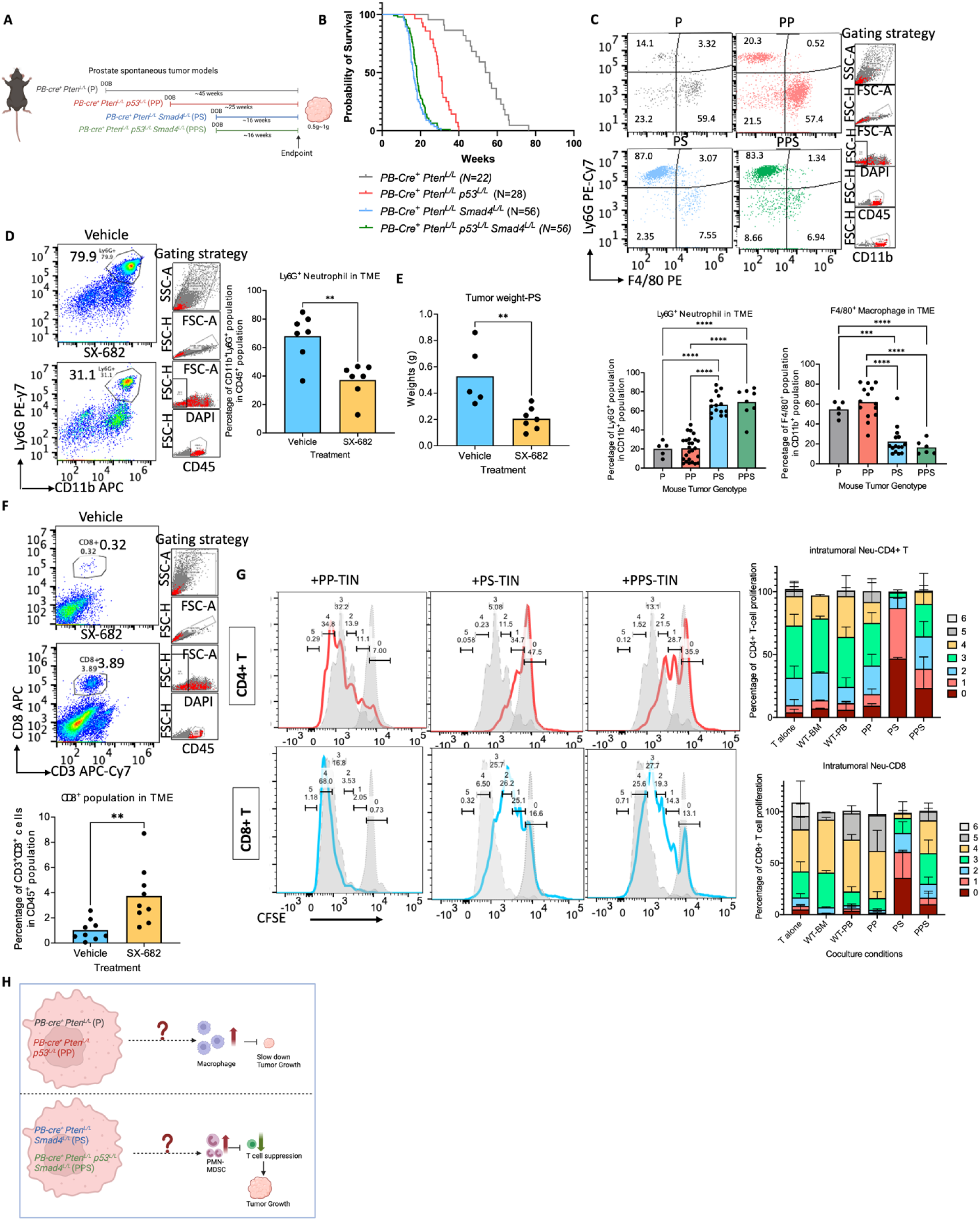
A distinct myeloid cell distribution across different genotypes of tumor models. (A)Illustration for mouse models used in the study. (B) Kaplan-Meier plot of the survival analysis of four representative tumor models. (C) FACS analyses showing dichotomous infiltration of Ly6G^+^F4/80^−^ cells(neutrophils) and Ly6G−F480+ cells (macrophages) in four representative tumor models. Plots are gated on CD45^+^ CD11b+ cells. The experiments were repeated at least five times with similar results. (D) FACS analyses and quantification showing TIN percentage in the tumor microenvironment of vehicle- and SX-682-treated tumor-bearing mice. Plots are gated on CD45^+^ cells. (E) The bar plot of PS tumor weights of vehicle- and SX-682-treated tumor-bearing mice. (F) FACS analyses and quantification showing the percentage of CD3^+^CD8^+^T cells in the tumor microenvironment of vehicle- and SX-682-treated tumor-bearing mice. Plots are gated on CD45^+^ cells. (G) FACS analyses of T cell CFSE assay, showing the T cell proliferation after coculturing with TINs purified from PP, PS, PPS tumors. (H) Illustration of the central question: how tumor-intrinsic factors shape NES and MES TMEs to influence tumor progression. Statistical significance: ns p > 0.05, * p < 0.05, ** p < 0.01, *** p < 0.001, **** p < 0.0001, Unpaired nonparametric Mann Whitney test.

In the four tumor genotypes, myeloid cell (CD11b^+^) populations occupied the major cell population in the immune cell (CD45^+^) populations, and the enrichment levels in the four genotypes were not significantly different (Figure S1B). Among the myeloid cell populations, tumor-infiltrating neutrophils (TINs) and macrophages (TIMs) were the most frequent and variable cell types, as confirmed by flow cytometry staining of Ly6G (neutrophil marker) and F4/80 (macrophage marker). TINs, defined as CD45^+^ CD11b^+^ Ly6G^+^, showed a predominated pattern within the microenvironment of both PS and PPS tumors. Conversely, an increase in TIMs, defined as CD45^+^ CD11b^+^ F4/80^+^, was observed in P and PP tumors (Figure 1C). Thus, we characterized PS and PPS tumors as neutrophil-enriched-subtype (NES) tumors (NES) and P and PP tumors as macrophage-enriched-subtype (MES) tumors. Notably, the NES tumor models demonstrated more aggressive tumor behavior, as evidenced by shorter survival periods (Figure 1B).

Previous research has reported that NES tumors were consistently immunosuppressive and resistant to immune checkpoint blockade (ICB) treatment, while macrophage-enriched tumors were responsive to ICB treatment in triple-negative breast cancer (TNBC)^45^. To assess whether TINs and TIMs exhibit pro-tumor or anti-tumor functions in our models, we pharmacologically depleted these cells *in vivo*. Neutrophils were depleted using the potent CXCR2 inhibitor SX-682^24^. After oral-garaging SX-682, the percentage of TINs significantly decreased (Figure 1D). Blocking TINs in PS tumors significantly reduced tumor progression in PS models (Figure 1E), accompanied by an increased proportion of CD8^+^ T cells (Figure 1F). In contrast, this *in-vivo* depletion of TINs did not affect tumor progression in PP models, suggesting that neutrophils in PP tumor microenvironment did not have pro-tumoral functions (Figure S1C). Other than TIN depletion, TIMs were depleted using a CCR2 inhibitor (CCR2i). After applying CCR2i, the percentage of TIMs decreased significantly (Figure S1D); however, no reduction in PP tumor weights was observed; instead, the trend indicated a potentially anti-tumoral role for macrophages in PP tumors (Figure S1E).

Functional CFSE assays confirmed the immunosuppressive capacity of TINs derived from PS and PPS tumors, which significantly inhibited CD3 and CD28 antibody-stimulated T-cell proliferation. However, neutrophils isolated from PP tumors were not immunosuppressive (Figure 1G). This immunosuppressive activity of TINs in PS and PPS tumors suggested the characteristic of polymorphonuclear myeloid-derived suppressor cells (PMN-MDSCs). This was also confirmed by the high expression of MDSC-associated markers, such as *Arg1* and *Nox2*, in neutrophils isolated from PS and PPS tumors. (Figure S1F).

Therefore, prostate tumors from different genotypes exhibited distinct immune cell accumulation in the TME and can be grouped into NES tumors and MES tumors. NES tumors displayed high immunosuppression in the tumor microenvironment and their tumor bearing mice performed poorer survivals, compared to MES tumors.

### Tumor-Intrinsic Factors Contribute to Immunosuppression in NES Tumor Microenvironment (TME)

Tumor-intrinsic factors contribute significantly to immunosuppression within the tumor microenvironment (TME). In this study, we aim to identify the genetic tumor-intrinsic factors that influence the formation of NES- or MES-type TMEs, as illustrated in (Figure 1H). To investigate the mechanism behind the high immunosuppressive neutrophil enrichment in NES tumors, we first analyzed an online dataset (GSE25140), the microarray of the same mouse models used in our study. They included two MES models P and PP prostate tumors and one NES model PS prostate tumor^21^. We identified 114 Gene Ontology (GO) signaling pathways enriched in PS(NES) vs. PP(MES) and 155 pathways enriched in PS(NES) vs. P(MES), with 21 overlapping pathways (Figure S2A). Notably, pathways related to neutrophil migration and chemokine activity were particularly enriched among these overlapping pathways. *Cxcl1, Cxcl2, Cxcl3, Cxcl5*, and *Ccl20* were found at higher levels in PS(NES) tumors compared to P and PP(MES) tumors (Figure S2B). These cytokines are all neutrophil-recruiting chemokines and indicated the high neutrophil chemotaxis in PS(NES) tumors. Besides, this chemotaxis activity was confirmed by *in-vivo* transplantation experiments. We injected mTomato^+^ neutrophils into tumor-bearing mice of different models and found an enriched neutrophil accumulation in PS(NES) tumors compared to PP(MES) counterparts (Figure S2C), suggesting that PS(NES) tumors were more capable of attracting neutrophils by expressing higher levels of neutrophil-recruiting chemokines.

To elucidate the cellular origins and signaling molecules governing the immunosuppressive TME within NES tumor cells, we used tumor cell lines from NES tumors (PS, PPS) and MES tumors (P, PP). Specifically, *Pten*-cap8 is derived from P prostate tumors (MES), 6240-PP and 7068-PP are isolated from PP prostate tumors (MES), TS3132, 06x-1, and 522665 are derived from PS prostate tumors (NES), and 9236 and 9320 are from PPS prostate tumors (NES) (Figure 2A). *Smad4* loss was confirmed by western blotting in these established tumor cell lines from each genotype (Figure S2D). Transwell migration assays demonstrated that PS tumor cells have a higher capacity of neutrophil recruitment, which aligns with the *in-vivo* observations (Figure 2B). To explore the cellular determinants of TIN accumulation, we collected conditioned medium and performed a cytokine array. The results revealed that PS tumors secreted higher levels of cytokines and chemokines, potent mediators of neutrophil recruitment such as CXCL1 and CXCL2 (Figure 2C). PP conditioned media had higher levels of CCL2, consistent with their higher macrophage levels (Figure 2C).

**Figure 2:**
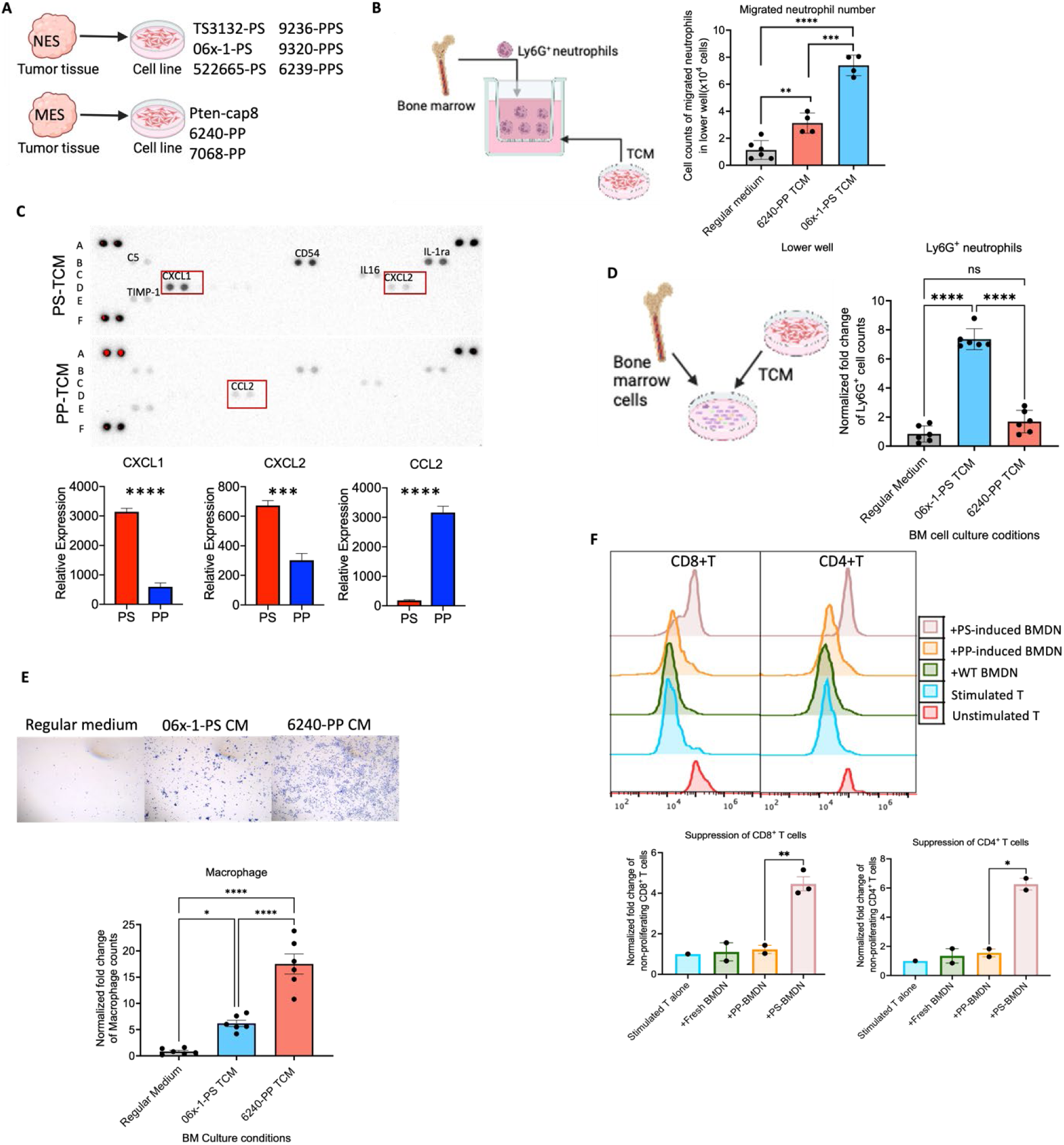
Tumour-intrinsic factors contribute to immunosuppression in TME. (A) Illustration for derived tumor cell lines. (B) Bar plot of the cell count number of Ly6G^+^ neutrophils attracted into the lower well of tumor conditional medium. (C) Bar plot of the cell count number of Ly6G^+^ neutrophils induced from bone marrow cells by PS conditional medium and PP conditional medium. (D) Crystal violet staining and quantification of attached cells from induced bone marrow cells by PS conditional medium and PP conditional medium. (E) The flow cytometry analysis of T cell CFSE assay, showing the T cell proliferation after coculturing with bone marrow-derived neutrophils by PP or PS conditional medium. (F) The cytokine array of PP and PS conditional medium. Statistical significance: ns p > 0.05, * p < 0.05, ** p < 0.01, *** p < 0.001, **** p < 0.0001, Unpaired nonparametric Mann Whitney test.

Besides the chemotaxis activity, we also checked whether NES tumors and MES tumors can influence on the induction and immunosuppressive activities of neutrophils/ macrophages. Using conditioned medium from PP and PS cell lines to induce wild-type bone marrow neutrophils, we demonstrated that PS-conditioned medium induced a higher number of neutrophils (Figure 2D), while PP-conditioned medium induced a higher number of macrophages (Figure 2E). Functional assays confirmed the immunosuppressive capacity of PS-conditioned medium-induced neutrophils, which suppressed both CD8^+^ and CD4^+^ T cells, highlighting the immunosuppressive induction by PS(NES) tumor cells (Figure 2F). These results suggested the effect of PS(NES) tumor cells on the differentiation and immunosuppression of neutrophils.

These findings underscore certain tumor-intrinsic factors in influencing the differentiation and activation, and recruitment of myeloid cells and in shaping the immune landscape of the TME.

### Protein-Arginine Deiminase (PADI) Activity Upregulated in PS and PPS Tumors and Its Role in Immunosuppressive Neutrophil Accumulation

To gain a comprehensive understanding of the mechanisms underlying the immunophenotypes associated with NES and MES tumors, we began by examining MES models. We observed that *Ccl2* was upregulated in PP cell lines (Figure 2C) and is known to function as a macrophage-recruiting chemokine^9^. Given the involvement of TGF-β-Smad4 signaling, we overexpressed *Smad4* in a PS cell line and observed that *Ccl2* expression was restored (Figure S2E). Analysis of prostate cancer transcriptomic data from the Prostate Cancer Transcriptomics Atlas revealed a positive correlation between *Ccl2* expression and the TGF-β signaling signature (Figure S2F), highlighting the role of TGF-β signaling in macrophage recruitment and the establishment of a MES-type TME.

To investigate NES tumors, we performed bulk RNA-seq analysis on tumor cell lines representing MES (P, PP) and NES (PS, PPS) phenotypes. Differentially expressed genes (DEGs) were identified, revealing that P and PP tumor cells exhibited similar expression patterns, while PS and PPS tumor cells showed distinct but similar expression patterns to each other. In the comparison between PS and PP cells, 1,418 genes were upregulated (logFC(PS vs. PP) > 1, P value < 0.05, FDR < 0.05), whereas 1,381 genes were downregulated (logFC(PS vs. PP) < -1, P value < 0.05, FDR < 0.05) (Figure 3A). Chemokine expression analysis across different tumor genotypes revealed elevated levels of CCL20, CXCL5, and CXCL2 in PS/PPS tumors compared to P/PP tumors, consistent with the microarray data from GSE25140 (Figure 3B).

**Figure 3:**
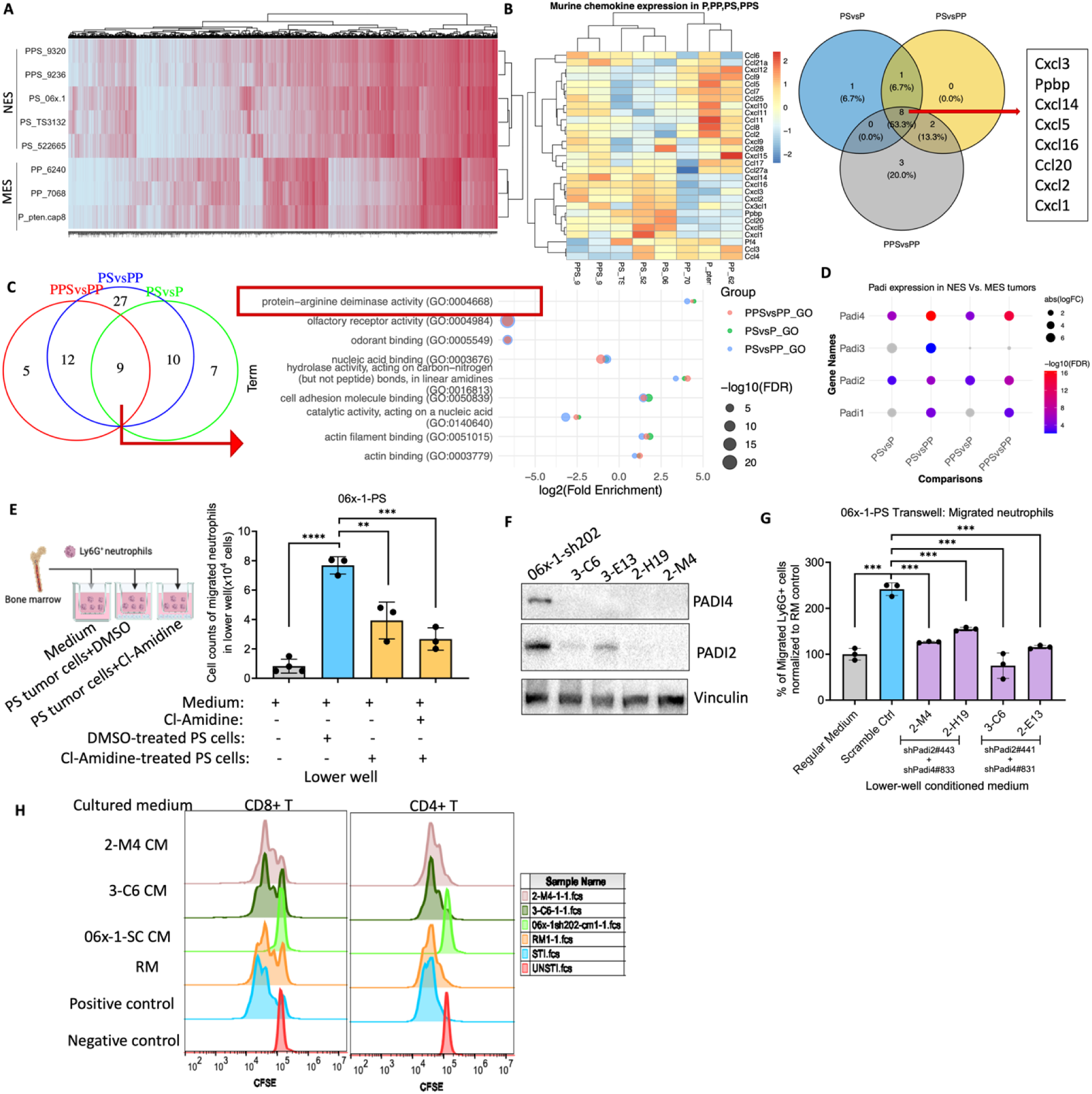
Protein-arginine deiminase (PADI) activity upregulated in PS and PPS tumors and its role in immunosuppressive neutrophil accumulation. (A) Heatmap of RNAseq data of P, PP, PS, PPS tumor cell lines. (B) Volcano plot of Chemokine profiling for four tumor genotypes. (C) Venn plot of Gene ontology molecular function enrichment analysis of PS vs. P, PS vs. PP, PPS vs. PP comparison(left) and dot plot showing the overlapped 9 signaling pathways (right). (D) Heatmap of the RNA levels of Padi genes in P, PP, PS, PPS cells. (E) Transwell assay of migrated neutrophil count in vehicle- or Cl-Amidine-treated 06x-1-PS. (F) Western blotting showing knockdown of Padi2 and Padi4. (G) Transwell assay of migrated neutrophil count in scramble control or shPadi2/4 KD of 06x-1-PS. (H) CFSE assay of BMDN induced by PS-scramble ctrl CM or PS-shPadi2/4KD CM or regular medium. Statistical significance: ns p > 0.05, * p < 0.05, ** p < 0.01, *** p < 0.001, **** p < 0.0001, Unpaired nonparametric Mann Whitney test.

GO_Molecular Function (GO_MF) analysis of comparisons between NES and MES tumor cells (PS vs. P, PS vs. PP, and PPS vs. PP cells) revealed nine overlapping pathways, with protein-arginine deiminase activity (GO:0004668) being the most enriched in NES (PS and PPS) cells compared to MES (P and PP) cell lines (Figure 3C). Citrullination, induced by Protein-arginine deiminases (PADIs), converts arginine to citrulline. By comparing the NES (PS and PPS) tumors and MES (P and PP) tumors, the expression levels of *Padi4* and *Padi2* are consistently and significantly higher in NES tumors compared to MES cell lines (Figure 3D). Western blot analysis of tumor cells isolated directly from tumor tissues confirmed higher levels of PADI4 and PADI2 in in PS models compared to PP models (Figure S3A). The expression of PADI2 and PADI2 also showed in other murine prostate tumor cell lines (Figure S3B). Cell fragmentation western blot analysis confirmed the localization of PADI1 in cells (Figure S3C). Immunohistochemistry (IHC) revealed nuclear localization of PADI4 in PS tumors, while nuclear PADI4 was undetectable in PP tumors (Figure S3D)

To investigate whether PADI2/4 contribute to the high levels of neutrophil accumulation in PS tumors, we treated PS cells with the pan-PADI inhibitor Cl-Amidine. This treatment reduced the neutrophil-attracting ability of PS cells (Figure 3E, Figure S3E) and significantly decreased chemokine expression, *Cxcl5*, *Ccl20*, and *Cxcl2*, levels in treated PS/PPS cells (Figure S3F-H). shRNA-mediated knockdown of *Padi2* and *Padi4* in the PS cell line, 06x-1 and 522665(Figure 3F) decreased the activity to recruit neutrophils, compared to the scramble control PS cell lines (Figure 3G, Figure S3I). The knockdown also declined the expression of the expression of chemokines *Cxcl5*, *Ccl20*, and *Cxcl2* in PS cells (Figure S3J). Conversely, overexpression of *Padi2* and/or *Padi4* in PP cell lines led to a dramatic increase in chemokine expression (Figure S3K-L), and rescuing *Padi2*/*Padi4* expression in knockdown cells(2-M4) restored certain neutrophil-recruiting chemokine expressions, such as *Cxcl5*, *Ccl20* (Figure S3M).

In addition to the effect on the chemotaxis activities, shRNA knockdown of *Padi2* and *Padi4* in the PS cell line also changed the ability to induce immunosuppressive neutrophils. We collected the tumor conditioned medium to culture naive bone marrow cells and isolated Ly6G^+^ neutrophils for the T cell CFSE assay. The induced neutrophils from Padi2/Padi4 double knockdown PS tumors (2-M4, 3-C6) were unable to suppress CD8^+^ and CD4^+^ T cells, compared to the induced neutrophil from scramble control PS tumor cells (Figure 3H), indicating the PADI2 and PADI4 were essential for PS tumor cells to induce immunosuppression.

Our findings indicate that PADI2 and PADI4 enriched in NES tumor cells are crucial for both neutrophil recruitment and immunosuppressive neutrophil induction in the NES TME. These enzymes play a key role in regulating cytokine expression that contributes to the immunosuppressive environment in NES (PS and PPS) tumors.

### Enhanced Histone Citrullination Upregulates Cytokine Expression to Promote Immunosuppressive Neutrophils

To investigate how the knockdown of *Padi2* and *Padi4* affects the neutrophil recruitment capacity of tumor cells, we performed a conditioned medium array of PS-scramble control cells and PS-sh*Padi2/4* cells and found that the expression level of TNFα, CXCL2 and GM-CSF decreased significantly after the suppression of *Padi2/4* (Figure 4C). Furthermore, we performed an RNA-seq to compare PS-scramble control cells with PS-sh*Padi2/4*(2-M4) cells, as well as vehicle and Cl-Amidine-treated PS cells. For the top extracellular genes that both were negatively regulated after the inhibition of Padi2/4 genetically and pharmalogically, we identified 197 overlapping genes. Among these, *Tnf*, *Ccl20*, and *Cxcl5* stood out as chemokines that regulate neutrophil recruitment and function, consistent with our qRT-PCR and cytokine assay results (Figure 4D). Gene Ontology (GO) analysis highlighted the key primary pathways that potentially influence PMN-MDSC accumulation (Figure 4E).

**Figure 4:**
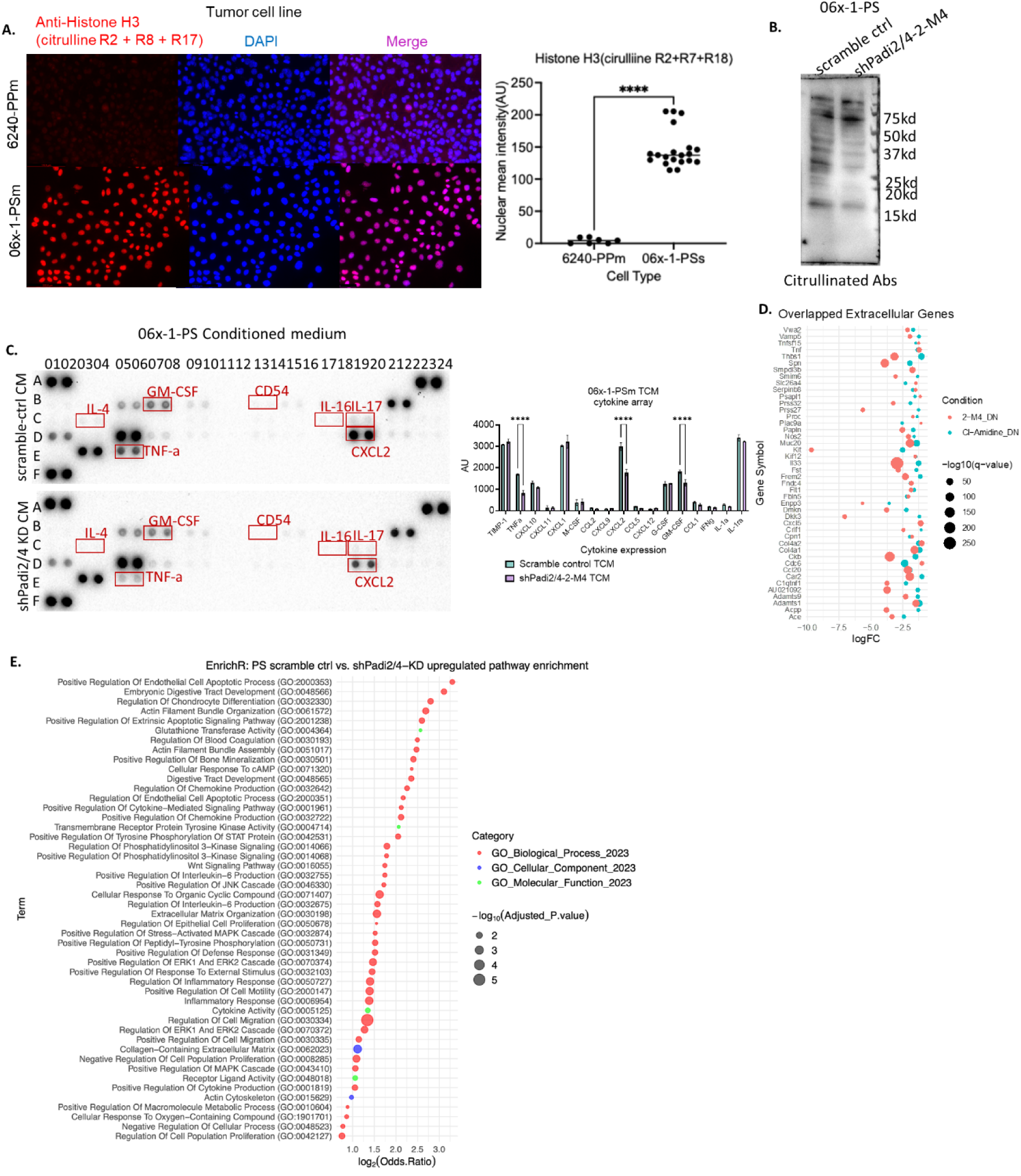
Enhanced histone citrullination promotes cytokine expression, leading to the recruitment of immunosuppressive neutrophils. (A) Immunocytochemistry analysis and quantification of Histone H3 (citrulline R2+R8+R17) expression and localization in 6240-PP and 06x-1-PS cells. (B) Western blot analysis of citrullinated proteins in scramble control and shPadi2/4 knockdown (KD) 06x-1-PS cells. (C) Cytokine array analysis and quantification of conditioned medium from scramble control and shPadi2/4 KD 06x-1-PS cells. (D) Top-enriched extracellular genes consistently downregulated in shPadi2/4 KD 06x-1-PS cells compared to scramble control, and in Cl-Amidine-treated 06x-1-PS cells compared to vehicle-treated cells. (E) Top enriched Gene Ontology (GO) terms for genes co-upregulated in scramble control 06x-1-PS cells compared to shPadi2/4 KD 06x-1-PS cells (adjusted p < 0.05), identified using EnrichR. Statistical significance: ns p > 0.05, * p < 0.05, ** p < 0.01, *** p < 0.001, **** p < 0.0001, Unpaired nonparametric Mann Whitney test.

PADI enzymes mediate citrullination, a post-translational modification that can significantly influence gene transcription^28,29^. Among the PADI family, PADI2 and PADI4 are known to induce histone citrullination, which affects chromatin structure and gene expression, contributing to tumor progression^46–48^. Previous studies have shown that PADI4 were able to target multiple sites in histones H3 and H4 and PADI2-mediated citrullination of H3R26 facilitates transcriptional activation driven by estrogen or androgen ^35,46,48^. Using immunofluorescent analysis, we observed high levels of histone citrullination (R2+R7+R18) in 06x-1-PS cells, compared to 6240-PP cells (Figure 4A), and in 522665-PS cells, compared to 6240-PP cells (Figure S4A). We also checked the histone citrullination level in tumor tissues and found that 638972-PS cyrosection tissue had higher histone citrullination(R2+R8+R17) signal, compared to 620706-PP cyrosection tissue (Figure S4B). In addition to histone citrullination (R2+R8+R17), the total protein citrullination level were performed by Western blot analysis and further demonstrated a decrease in total citrullinated proteins at certain sizes after *Padi2/4* knockdown (KD) (Figure 4B). Furthermore, we examined different histone arginine sites of citrullination. After shPadi2/4 knockdown or treatment with pan-PADI inhibitor Cl-Amidine, we found a reduction in citrullinated Histone H3 at R2+R8+R17 and R26 following Ca2+ treatment (Figure S4C).

Histone citrullination regulates gene expression by modifying chromatin accessibility. We performed ATAC-seq on scramble control and sh*Padi2/4* KD 06x-1-PS cells to examine chromatin openness. Analysis of chemokines upregulated in NES (PS and PPS) tumor cells versus MES (PP) cells (Figure 3B) revealed *Ccl20*, *Ppbp*, *Cxcl1*, and *Cxcl2* as top-enriched genes (Figure S4D). We then selected genes showing significantly higher chromatin openness in the scramble control group (logFC>0.5, p<0.05) and cross-referenced them with upregulated genes from the RNA-seq, identifying 571 overlapping genes (Figure S4E).

Our findings reveal that enhanced histone citrullination, driven by increased PADI2/4 activity, activated key signaling pathways and upregulated neutrophil-related cytokine expression in tumor cells. This promotes the recruitment and accumulation of immunosuppressive neutrophils within the tumor microenvironment, thus facilitating immune evasion.

### Smad4 Ablation Upregulates PADI2/4 Expression via YAP Signaling

We next investigated the mechanistic drivers regulating the induction of PADI2 and PADI4 in NES (PS and PPS) tumors. Notably, in our models, NES tumors all exhibited *Smad4* loss. Clinical data from the TCGA dataset revealed that SMAD4 expression is significantly downregulated in primary tumors (n=497) compared to normal tissue (n=52) (Figure S4F). Patients with lower SMAD4 expression in prostate adenocarcinoma have poorer survival outcomes compared to those with higher SMAD4 expression (Figure S4G). Furthermore, as prostate cancer progresses and the Gleason score increases, the expression of SMAD4 and the enrichment of TGFβ signaling show a decreasing trend (Figure S4H-I).

Yes-Associated Protein 1 (YAP1), previously reported to be hyperactivated in *Smad4*-null prostate tumors ^23^, was found to be activated in PS cells compared to PP cells (Figure 5A). Similar trends were observed in tumor tissues (Figure S5A). Inhibition of YAP1 by YAP inhibitor, verteporfin, in PS tumor cells decreased *Padi2* and *Padi4* expression compared to the vehicle control (Figure 5B). The decrease in expression was also observed in neutrophil-recruiting chemokines *Cxcl5* and *Ccl20* (Figure S5B). Ablation of *Yap1* by shRNA knockdown or CRISPR-Cas9 knockout downregulated *Padi2* and *Padi4* expression in PPS cells (Figure 5C, Figure S5C-D). Additionally, Tead4 knockdown also resulted in decreased expression of *Padi2* and *Padi4* (Figure S5E-F). The YAP1 phosphorylation mutant (YAP-S127A), which activates YAP signaling (Figure S5G-H), induced *Padi2* and *Padi4* expression in TRAMP-C1 and TRAMP-C2 tumor cells (Figure 5D) and also enhanced neutrophil-recruiting chemokines *Cxcl5* and *Ccl20* expression (Figure S5I-J). Notably, the YAP/TAZ target signature was positively correlated with augmented *Padi2* and *Padi4* mRNA levels in human prostate tumors in the TCGA dataset (Figure 5E).

**Figure 5:**
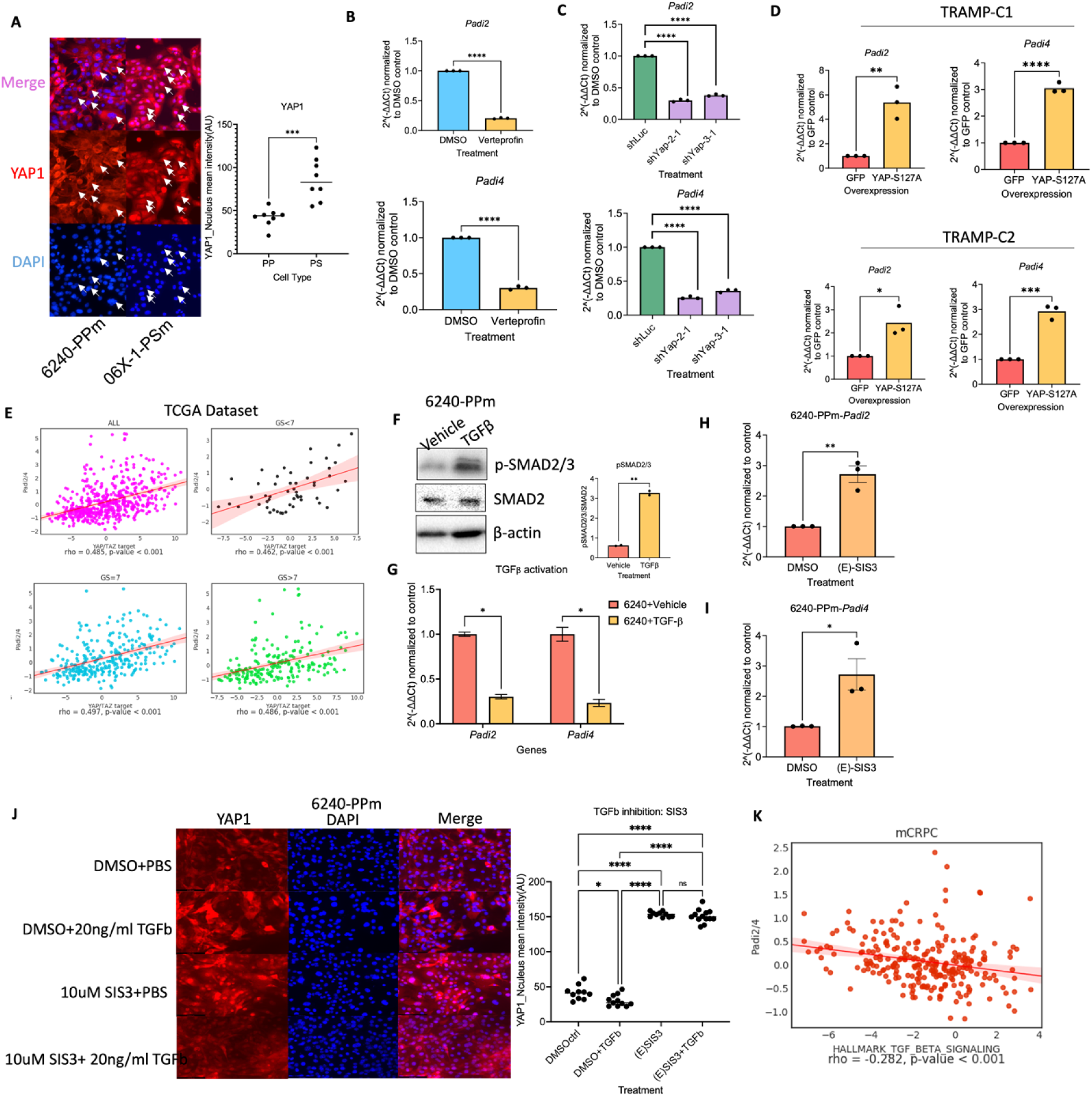
Smad4 ablation upregulates PADI2/4 expression via YAP signaling. (A) Immunocytochemistry analysis of YAP1 expression and localization in 6240-PP and 06x-1-PS cells. (B) qPCR analysis of Padi2 and Padi4 expression in verteporfin-treated versus DMSO-treated 06x-1-PS cells. (C) qPCR analysis of Padi2 and Padi4 expression in shLuc- and shYap1-KD 6239-PPS cells. (D) qPCR analysis of Padi2 and Padi4 expression in GFP- and YAP-S127A-expressing TRAMP-C1 and TRAMP-C2 cells. (E) Correlation between Padi2/4 expression and the YAP/TAZ target signature in the TCGA dataset, assessed using the PCTA web tool. (F) Western blot analysis of phosphorylated SMAD2/3 (pSMAD2/3) and total SMAD2 in vehicle- and 10 ng/ml TGFβ-treated 6240-PP cells, with β-actin as the loading control. (G) qPCR analysis of Padi2 and Padi4 expression in vehicle- and 10 ng/ml TGFβ-treated 6240-PP cells. (H) Western blot analysis of phosphorylated SMAD2/3 (pSMAD2/3) and total SMAD2 in vehicle- and 10 μM (E)SIS3-treated 6240-PP cells, with vinculin as the loading control. (I) qPCR analysis of Padi2 and Padi4 expression in vehicle- and 10 μM (E)SIS3-treated 6240-PP cells. (J) Immunocytochemistry analysis of YAP1 expression and localization in cells treated with DMSO+PBS, DMSO+10 ng/ml TGFβ, 10 μM (E)SIS3+PBS, and 10 μM (E)SIS3+10 ng/ml TGFβ. (K) Correlation between Padi2/4 expression and the YAP/TAZ target signature in the Prostate Cancer Transcriptome Atlas (PCTA) dataset, assessed using the PCTA web tool. Statistical significance: ns p > 0.05, * p < 0.05, ** p < 0.01, *** p < 0.001, **** p < 0.0001, Unpaired nonparametric Mann Whitney test.

To study the genotypic background of NES tumors, we investigated how *Smad4* loss induced YAP-PADI2/4-NES phenotype. Since SMAD4 is an essential element in the TGFβ signaling pathway, we used recombinant TGFβ to treat PP cells to stimulate TGFβ signaling activation (Figure 5F). After the activation of TGFβ signaling, the expression of *Padi2* and *Padi4* decreased (Figure 5G). When we used a potent TGFβ signaling inhibitor, (E)-SIS3, to treat PP cells, the expression of *Padi2* and *Padi4* increased (Figure 5H-I). We also found that YAP1 signaling increased in the nucleus after (E)-SIS3 treatment (Figure 5J). We then examined the relationship between TGFβ signaling signature and *Padi2/4* levels in the clinical patient samples of the Prostate Cancer Transcriptomics Atlas (PCTA) datasets. In a group of CRPC patients, Padi2/4 levels were negatively correlated with the TGFβ signaling signature (Figure 5K).

Collectively, these results suggest that TGFβ signaling pathways regulated YAP signaling and thus the expression *Padi2* and *Padi4* are enhanced in PS and PPS tumor cells via the activation of YAP1 signaling. This enhancement contributes to the immunosuppressive neutrophil accumulation in the TME.

### Inhibition of PADI Activity Suppresses Neutrophil Accumulation in Prostate Tumors and Enhances ICB Efficacy

Given that *Padi2/4* ablation in PS cells abolished neutrophil migration *in vitro* (Figure 3G) and TINs in PS tumor models had protumoral functions (Figure 1E), we further investigated its effects *in vivo*. C57BL/6 mice were subcutaneously injected with either 06x-1-PS-scramble control or PS-sh*Padi2/4*-2-M4 cells. Knockdown of *Padi2/4* led to a 25-50% reduction in PS tumor progression compared to the scramble control (Figure 6A). However, in immunocompromised NOD-SCID mice, no differences in tumor growth were observed between the two groups (Figure 6B), suggesting that lymphocytes play a crucial role in the inhibition of tumor growth in sh*Padi2/4* knockdown tumors.

**Figure 6:**
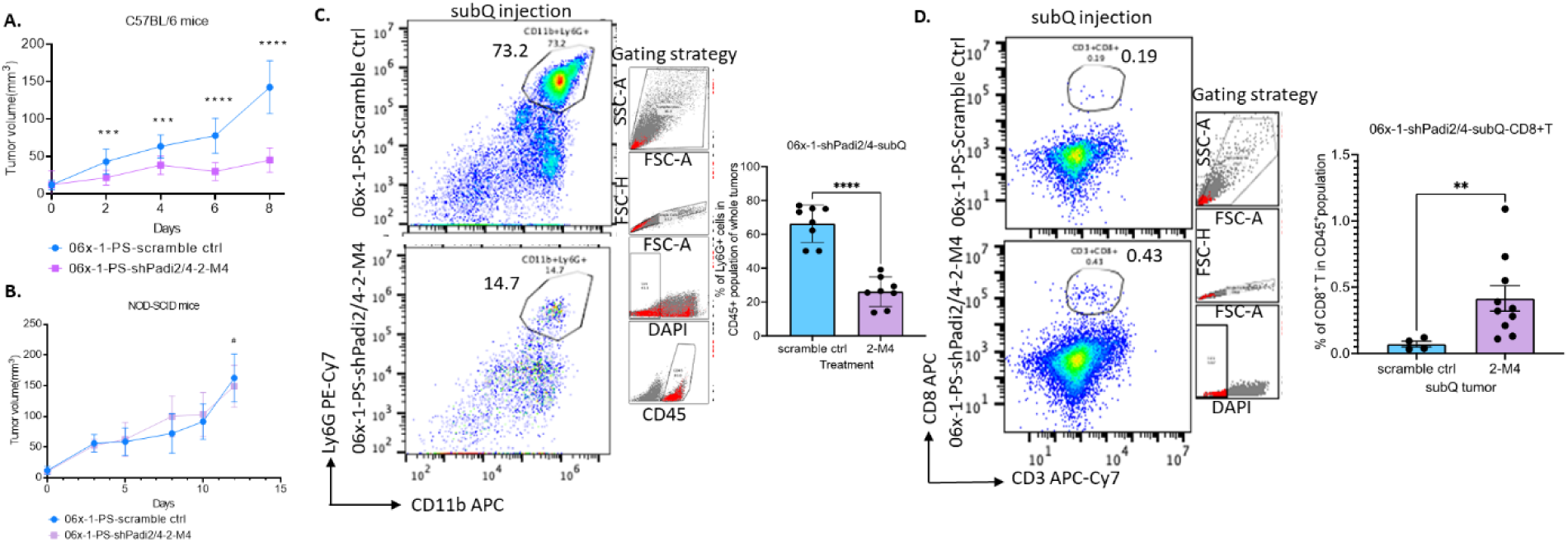
PADI knockdown suppresses neutrophil accumulation in prostate tumors. (A) Tumor growth curves of shScramble control and shPadi2/4 KD subcutaneous tumors in C57BL/6 mice, presented as tumor volume (mm^3^). (B) Tumor growth curves of shScramble control and shPadi2/4 KD subcutaneous tumors in NOD-SCID mice, shown as tumor volume (mm^3^). (C) FACS analysis and quantification of CD11b+ Ly6G+ tumor-infiltrating neutrophils (TINs) in the tumor microenvironment of shScramble control and shPadi2/4 KD subcutaneous tumors in C57BL/6 mice. (D) FACS analysis and quantification of CD3+ CD8+ T cell percentages in the tumor microenvironment of shScramble control and shPadi2/4 KD subcutaneous tumors in C57BL/6 mice. Statistical significance: ns p > 0.05, * p < 0.05, ** p < 0.01, *** p < 0.001, **** p < 0.0001, Unpaired nonparametric Mann Whitney test.

Next, we assessed the percentage of tumor-infiltrating neutrophils (TINs) in *Padi2/4* knockdown tumors, finding a significant decrease in TINs (Figure 6C), alongside a significant increase in CD8^+^ T cells (Figure 6D). In addition to the subcutaneous model, we established prostate orthotopic models and observed decreased neutrophil infiltration along with increased populations of both CD4^+^ and CD8^+^ T cells (Figure S6A-C).

In summary, these results demonstrate that inhibition of PADI activity, when combined with ICB treatment, can effectively suppress tumor progression, highlighting its potential as a therapeutic strategy for enhancing the efficacy of immunotherapy in prostate cancer.

### PADI2/4 Is Upregulated in Human Prostate Cancer and Correlated with PMN-MDSC Recruitment

Lastly, we want to reveal whether PADI2 and PADI4 are upregulated in human prostate cancer tissues and correlated with PMN-MDSC infiltration in the clinical samples. It has been reported that the levels of citrullinated proteins are higher in CRPC patients ^35^. To determine whether PADI2 and PADI4 are overexpressed in human prostate cancer, we first tested their expression in human prostate cancer cell lines and found that 22Rv1 and NCI-H660 have high levels of both PADI2 and PADI4 (Figure 7A). Additionally, these cell lines exhibited higher levels of CXCL5 expression compared to other cell lines (Figure 7B). We then analyzed RNAseq data from prostate cancer patients in the GSE21036 dataset ^49^ and found that both *PADI2* and *PADI4* are expressed at higher levels in tumor tissue than in normal prostate tissue (Figure 7C), with a tendency to increase as prostate cancer progresses (Figure 7D).

**Figure 7:**
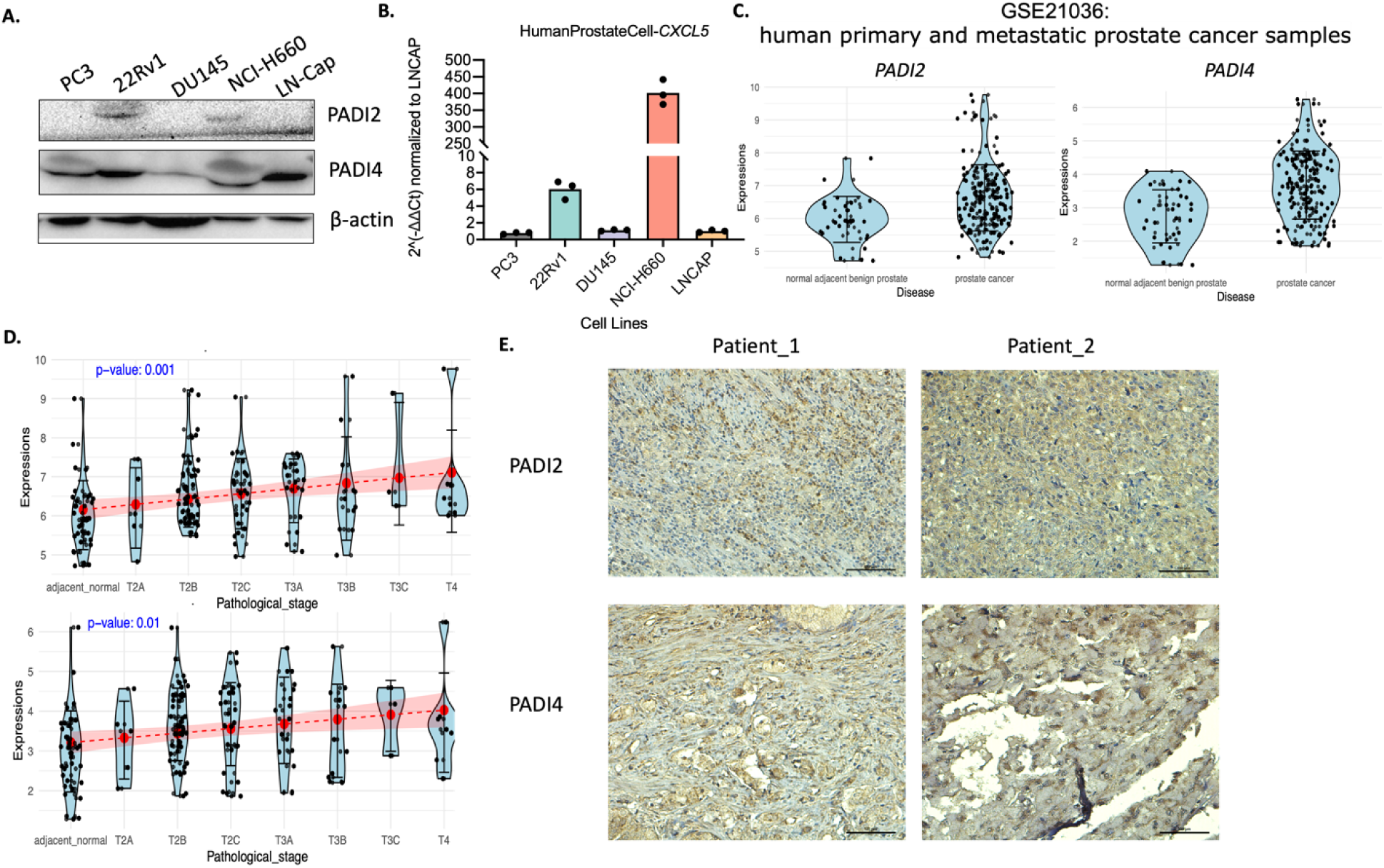
PADI2/4 is upregulated in human prostate cancer and correlates with PMN-MDSC recruitment. (A) Western blot analysis of PADI2 and PADI4 expression in human prostate cancer cell lines: PC3, 22Rv1, DU145, NCI-H660, and LN-Cap. β-actin served as the loading control. (B) qPCR analysis of Cxcl5 chemokine expression in human prostate cancer cell lines: PC3, 22Rv1, DU145, NCI-H660, and LN-Cap. (C) Normalized expression levels of PADI2 and PADI4 in normal adjacent benign prostate tissue and prostate cancer tissue (GSE21036 dataset). (D) Normalized expression levels of PADI2 and PADI4 across different pathological stages of prostate cancer (GSE21036 dataset), with red dots indicating mean expression levels at each stage. (E) IHC analysis of PADI2 and PADI4 expression in prostate cancer tissues from two patients. Statistical significance: ns p > 0.05, * p < 0.05, ** p < 0.01, *** p < 0.001, **** p < 0.0001, Unpaired nonparametric Mann Whitney test.

Next, we performed IHC staining of a human prostate cancer tissue for PADI2, and PADI4 and showed the expression of PADI2 and PADI4 in the prostate cancer patient tumor samples (Figure 7E). Given the lack of validated antibodies for human MDSCs in TMA analysis, we utilized a neutrophil score and MDSC score from ssGSEA and 39-gene MDSC signature from a previous report^50^. Using MSKCC Prostate Cancer Genomics Data Portal and prostate RNA-sequencing data from the Prostate Cancer Transcriptome Atlas (PCTA), we tested the correlation with the neutrophil/MDSC/PMN-MDSC signature and found that PADI2 and PADI4 are significantly correlated with the neutrophil score and MDSC score (Figure 7SA) and PMN-MDSC signature (Figure 7SB).

These findings in human prostate tumors, which parallel our murine observations, suggest that enhanced PADI2/4 expression is integral to MDSC infiltration in both mouse and human prostate cancer. This highlights the translational value of our study and underscores the potential of targeting PADI2/4 to modulate the tumor microenvironment and improve therapeutic outcomes in prostate cancer.

## Discussion

This study provides significant insights into the mechanisms by which tumor genotype influences the immune landscape of prostate cancer, particularly focusing on the role of *Smad4* loss in driving immunosuppressive neutrophil accumulation in TME. Our findings demonstrate that *Smad4*-null tumors (PS and PPS) are neutrophil-enriched (NES) and exhibit more aggressive behavior compared to macrophage-enriched (MES) tumors in *Pten^pc-/-^*(P) and *Pten^pc-/-^ p53^pc-/-^* (PP) models. This distinction is crucial for understanding how different genetic backgrounds can dictate immune cell composition and tumor progression.

The hyperactivation of YAP1 signaling in *Smad4*-null tumors leads to the upregulation of PADI2 and PADI4, enzymes involved in citrullination, which in turn facilitate the recruitment of immunosuppressive tumor-infiltrating neutrophils (TINs). Furthermore, our data indicate that inhibiting YAP1 signaling through verteporfin, or genetic ablation significantly reduces PADI2/4 expression and subsequent neutrophil recruitment.

*In-vivo* experiments further validated the role of PADI2/4 in modulating the TME. Knockdown of *Padi2/4* in PS tumor cells led to a marked reduction in TIN accumulation, enhanced infiltration of CD8^+^ T cells, and partial tumor remission. We anticipate that combining *Padi2/4* knockdown with immune checkpoint blockade (ICB) therapy will result in complete tumor regression in mouse models, highlighting the potential of this combinatorial approach in enhancing immunotherapeutic efficacy.

Human prostate cancer data corroborated our murine findings, showing elevated PADI2/4 expression in tumor tissues compared to normal prostate tissues, and a positive correlation with neutrophil-attracting chemokines and a PMN-MDSC signature. These observations suggest that the mechanisms identified in our mouse models are relevant to human disease and underscore the translational potential of targeting PADI2/4 in prostate cancer therapy.

Our study also sheds light on the interplay between genetic loss of Smad4, TGFβ, and YAP1 signaling in regulating PADI2/4 expression and subsequent PMN-MDSC infiltration. While our findings elucidate key aspects of PADI2/4-mediated immunosuppression in the TME, further research is needed to explore the biochemistry of citrullination in greater depth. Understanding the precise biochemical mechanisms by which PADI2 and PADI4 modify proteins and how these modifications influence immune cell behavior could reveal additional therapeutic targets. Additionally, exploring the broader role of citrullination in other immune cell types within the TME and its impact on tumor progression could provide a more comprehensive understanding of its role in cancer biology.

Future studies should also investigate the potential side effects and optimal dosing strategies for PADI inhibitors in clinical settings. Combining PADI inhibitors with other immunotherapies or targeted treatments may enhance their efficacy, but the interactions between these therapies need careful evaluation.

## Acknowledgements

The authors thank all members of the lab for their support. We would like to acknowledge the following funding support: summer fellowships from Berthiaume Institute for Precision Health at University of Notre Dame (Y.Z.), National Institutes of Health grant R01CA248033 (X.L.), R01CA280097 (X.L.), R01CA297220 (XL), Department of Defense Congressionally Directed Medical Research Programs grants W81XWH2010332 (X.L.), HT94252310010 (X.L.) and HT94252310613 (X.L.), and Boler Family Foundation (X.L.) at University of Notre Dame.

## Author Contributions

Conceptualization, X.L.; methodology, Y.Z, and X.L; investigation, Y.Z, Y.L, and D.G; writing-– original draft, Y.Z; writing-–review & editing, Y.Z, and X.L.; supervision, X.L.

## Conflict of Interest

The authors declare no conflict of interest.

## Materials and Methods

### Mice Strains and Cell Culture

For mouse strains, *Pten^pc-/-^*, Pten *^pc-/-^ Smad4 ^pc-/-^*, *Pten ^pc-/-^ p53 ^pc-/-^* and *Pten ^pc-/-^ p53 ^pc-/-^ Smad4 ^pc-/-^* models were developed previously^21^. C57BL/6 WT strain was obtained from The Jackson Laboratory. Mice were maintained in pathogen-free conditions at the Freimann Life Science Center (FLSC). All manipulations were approved by IACUC at University of Notre Dame.

For cell lines, *Pten*-cap8 were obtained on ATCC (ATCC, CRL-3033, Manassas, VA, USA). Pten *^pc-/-^ Smad4 ^pc-/-^* prostate cell lines, which have been described previously^21,36^. PPS, a C57BL/6-syngeneic cell line isolated from prostate tumors of *Pten ^pc-/-^ p53 ^pc-/-^ Smad4 ^pc-/-^* mice. PP, a C57BL/6-syngeneic cell line isolated from prostate tumors of *Pten ^pc-/-^ p53 ^pc-/-^*mice.

Cells were cultured in Dulbecco’s Modified Eagle Medium (DMEM, VWR, 16750-074, Radnor, PA, USA) supplemented with 10% fetal bovine serum (FBS, HyClone, GE Healthcare Life Sciences SH30396.03, Marlborough, MA, USA) and 1X Penicillin/Streptomycin (Caisson Labs, PSL01, Smithfield, UT, USA). All cell lines tested for Mycoplasma were negative within 6 months of performing the experiments.

### Flow Cytometry

Cells (1 × 10^6^) were incubated with diluted anti-Mouse CD16/CD32 (Fc Shield, 1:500, Cytek Biosciences, 70-0161-U500, San Diego, CA, USA) for 10 min on ice. Cells were then washed and stained with primary antibodies for 30min on ice. Primary antibodies include anti-Mouse-CD45-FITC (Cytek Biosciences, 35-0451-U100, San Diego, CA, USA), anti-Mouse-CD11b-APC (Cytek Biosciences, 20-0112-U100, San Diego, CA, USA), anti-Mouse-Ly6G-PE-Cyanine7 (Cytek Biosciences, 60-1276-U100, San Diego, CA, USA), anti-Mouse-F4/80-PE (Cytek Biosciences, 50-4801-U100, San Diego, CA, USA), anti-Mouse-CD3-APC-Cyanine7 (Cytek Biosciences, 25-0032-U100, San Diego, CA, USA), anti-Mouse-CD8-APC (Cytek Biosciences, 20-0081-U100, San Diego, CA, USA), anti-Mouse-CD4-PE (Cytek Biosciences, 50-0041-U100, San Diego, CA, USA). Dead cells were distinguished by DAPI (0.5 μg/mL) staining. Flow cytometry analyses were performed on CytoFLEX S V4-B2-Y4-R3 Flow Cytometer (Beckman Coulter, C09766, Mississauga, Ontario, Canada). Quantification and data presentation were done with FlowJo Software version 10.4.

### Western Blot

Cells were lysed in 1x RIPA buffer, and the lysates were mixed with 5X SDS loading buffer. The samples were run on an SDS-PAGE gel and electrotransferred onto a 0.45μm nitrocellulose membrane using Trans-Blot® Turbo™ Transfer System (Bio-Rad, 1704150EDU, Hercules, CA, USA). Following the electrotransfer, the membrane was blocked with 5% milk in TBST at RT for 1h, then incubated with primary antibody diluted in 1x TBST at 4 °C overnight. The next day, the membrane was washed with 1x TBST and incubated with an HRP-conjugated secondary antibody in 5% milk in TBST at RT for 1h. After washing with 1x TBST, the membrane was developed using Clarity Western ECL Substrate (Bio-Rad, 1705061, Hercules, CA, USA) and imaged with a ChemiDoc Imaging System(Bio-Rad). The primary antibodies used included PADI1 (ABclonal, A10145, Woburn, MA, USA), PADI2 (Proteintech, 12110-1-AP, Rosemont, IL, USA), PADI4 (Proteintech, 17373-1-AP, Rosemont, IL, USA), Anti-Histone H3(citrulline R2 + R8 + R17) antibody (Abcam, ab281584, Cambridge, UK), Anti-Histone H3(citrulline R26) antibody (Abcam, ab212082, Cambridge, UK),, Anti-Citrulline antibody (Abcam, ab100932, Cambridge, UK), SMAD2 (Invitrogen, 51-1300, Waltham, MA USA), Phospho-SMAD2 (Ser465/467)/SMAD3 (Ser423/425) (Cell Signaling Technology, 8828T, Danvers, MA, USA), YAP (Cell Signaling Technology, 14074T, Danvers, MA, USA), Phospho-YAP (Ser127) (Cell Signaling Technology, 13008T, Danvers, MA, USA), Histone H3 (Cell Signaling Technology, 4499S, Danvers, MA, USA), β-actin (Santa Cruz Biotechnology, sc-47778, Dallas, TX, USA), GAPDH (Sigma-Aldrich, G9545, St. Louis, MO, USA), and Vinculin1(Millipore, 05-386, Burlington, MA, USA).

### Quantitative Reverse Transcription-Polymerase Chain Reaction (qRT-PCR)

Total RNA was isolated using a Total RNA Isolation Miniprep Kit (Bio Basic, BS1361, BUFFALO, NY, USA). Complementary DNA was synthesized using an All-In-One RT MasterMix (ABM, G490, Richmond, BC, Canada) according to the manufacturer’s instruction. Relative quantities of specific mRNA species were measured using SYBR Green qPCR Master Mix (Bimake, B21203, Houston, TX, USA) on CFX Connect Real-Time PCR System (Bio-Rad, Hercules, CA, USA). Amplification was performed with 40 cycles at 95°C for 15s and 60°C for 30s.

Data was analyzed with the 2^ΔΔCt method with the genes of interest normalized to Gapdh (housekeeping control). The primer sequences were as follows: *Nox2* (forward: ACC GGG TTT ATG ATA TTC CAC CT; reverse: GAT TTC GAC AGA CTG GCA AGA); *NOS2* (forward: TTC AGT ATC ACA ACC TCA GCA AG; reverse: TGG ACC TGC AAG TTA AAA TCC C); *Arg1*(forward: GTG GAA ACT TGC ATG GAC AAC; reverse: AAT CCT GGC ACA TCG GGA ATC); *Padi2* (forward: AAG GGG CTA TCC TGC TGG T; reverse: GAC CTT TTC GTC ACT ACA GTC); *Padi4* (forward: TCT GCT CCT AAG GGC TAC ACA; reverse: GTC CAG AGG CCA TTT GGA GG); *Cxcl5* (forward: GTT CCA TCT CGC CAT TCA TGC; reverse: GCG GCT ATG ACT GAG GAA GG); *Cxcl2* (forward: CCA ACC ACC AGG CTA CAG G; reverse: GCG TCA CAC TCA AGC TCT G); *Ccl20* (forward: GCC TCT CGT ACA TAC AGA CGC; reverse: CCA GTT CTG CTT TGG ATC AGC); *Tead4* (forward: CAA CCT GGA ACA TCC CAC GAT; reverse: GAA AGC CGA GAA CTC CAA CAT).

### T-cell Suppression Assay

The T-cell suppression assay was conducted following the procedure described previously ^23^. Ly6G^+^ neutrophils were isolated using magnetic bead sorting of MojoSort^TM^ Mouse Neutrophil Isolation Kit (BioLegend, 480058, San Diego, CA, USA), while CD3^+^ T cells were magnetically sorted using the MojoSort^TM^ Mouse CD3 T cell Isolation Kit (BioLegend, 480031, San Diego, CA, USA) and were labeled with 5μM CFSE division tracker (BioLegend, 1423801, San Diego, CA, USA). The assay was performed in 5μg/ml-anti-CD3 (BioLegend, 100302, San Diego, CA, USA)– coated 96-well plates. Isolated T cells were pre-treated with 2μg/ml anti-CD28 antibody (BioLegend, 102116, San Diego, CA, USA). For the coculturing, T-cell/neutrophil ratios were 0:1, 1:1, 1:2, and 1:4. After 72 hours, T-cell proliferation was assessed by flow cytometry, and suppression was determined by comparing the percentage of CFSE^+^ cells that divided in the presence of neutrophils to those that divided without neutrophils, as described by ^23^.

### Neutrophil Migration (Transwell) Assay

For the neutrophil migration (Transwell) assay, equal numbers of Ly6G^+^ neutrophils, isolated from C57BL/6 WT mouse bone marrow using the MojoSort^TM^ Mouse Neutrophil Isolation Kit (BioLegend, 480058, San Diego, CA, USA), were placed in the upper chamber of a transwell system (Costar Transwell, 6.5 mm, 8.0 μm, Tissue Culture Treated, Corning, 3464, Wilmington, NC, USA). Conditioned media from tumor cells under different conditions was added to the lower chamber, or tumor cells were pre-seeded on the bottom of the lower chamber. Cells were incubated for 24 hours at 37°C with 5% CO_2_, after which migrated cells in the bottom well were analyzed by flow cytometry. The absolute number of migrated Ly6G^+^ cells was determined by gating on these cells.

### RNA-sequencing

Raw counts were processed using the R package edgeR^37^. Lowly expressed genes were removed with filterByExpr, and library sizes were normalized using the trimmed mean of M-values. Dispersion was estimated, and a generalized linear model (GLM) was fitted with group as the design factor. Differential expression was assessed with likelihood ratio testing, and genes with *P* < 0.05 and |log_2_FC| > 1 were considered significant. The cell–cell communication was inferred with CellChat^38^ (mouse database) R package. Log-transformed expression data were used to identify overexpressed genes and ligand–receptor interactions, compute communication probabilities, and aggregate pathway-level networks.

### ATAC-seq Analysis

Paired-end ATAC-seq reads were aligned to the mm10 reference genome using Bowtie2^39^ (--very-sensitive, maximum fragment length 2000 bp), followed by sorting and indexing with Samtools^40^ Peaks were called with MACS2^41^(callpeak-f BAMPE), and replicate peak summits were concatenated, sorted, and merged with Bedtools^42^ to obtain a consensus peak set. Read counts per peak across all samples were quantified with bedtools multicov, generating a peak-by-sample count matrix. Consensus peaks were annotated relative to genomic features using ChIPseeker^43^ with the mm10 TxDband *org.Mm.eg.db*. Peak sequences were retrieved from the mm10 genome to calculate GC content, and GC bias was corrected using EDASeq^44^ within- and between-lane normalization. Normalized counts were analyzed with edgeR, where low-abundance peaks were filtered with filterByExpr, and a GLM likelihood ratio test was applied to identify differentially accessible regions. Peaks with *P* < 0.05 were considered significant and mapped back to genomic coordinates for downstream interpretation.

### Statistical Analysis

Data were displayed as mean ± standard deviation. Experiments were repeated at least three times with representative results shown. Comparisons between two groups were performed using the Unpaired nonparametric Mann Whitney test and Comparisons more than two groups were performed using one-way ANOVA with multiple comparisons by GraphPad Prism software v8.2 (GraphPad Software, San Diego, CA, USA). p < 0.05 was considered statistically significant.

## Notes

### Competing Interest Statement

The authors have declared no competing interest.

## References

(1) Umansky, V.; Sevko, A. Tumor Microenvironment and Myeloid-Derived Suppressor Cells. 10.1007/s12307-012-0126-7.

(2) Thorsson, V.; Gibbs, D. L.; Brown, S. D.; Wolf, D.; Bortone, D. S.; Ou Yang, T. H.; Porta-Pardo, E.; Gao, G. F.; Plaisier, C. L.; Eddy, J. A.; Ziv, E.; Culhane, A. C.; Paull, E. O.; Sivakumar, I. K. A.; Gentles, A. J.; Malhotra, R.; Farshidfar, F.; Colaprico, A.; Parker, J. S.; Mose, L. E.; Vo, N. S.; Liu, J.; Liu, Y.; Rader, J.; Dhankani, V.; Reynolds, S. M.; Bowlby, R.; Califano, A.; Cherniack, A. D.; Anastassiou, D.; Bedognetti, D.; Rao, A.; Chen, K.; Krasnitz, A.; Hu, H.; Malta, T. M.; Noushmehr, H.; Pedamallu, C. S.; Bullman, S.; Ojesina, A. I.; Lamb, A.; Zhou, W.; Shen, H.; Choueiri, T. K.; Weinstein, J. N.; Guinney, J.; Saltz, J.; Holt, R.; Rabkin, C. E.; Caesar-Johnson, S. J.; Demchok, J. A.; Felau, I.; Kasapi, M.; Ferguson, M. L.; Hutter, C. M.; Sofia, H. J.; Tarnuzzer, R.; Wang, Z.; Yang, L.; Zenklusen, J. C.; Zhang, J. (Julia); Chudamani, S.; Liu, J.; Lolla, L.; Naresh, R.; Pihl, T.; Sun, Q.; Wan, Y.; Wu, Y.; Cho, J.; DeFreitas, T.; Frazer, S.; Gehlenborg, N.; Getz, G.; Heiman, D. I.; Kim, J.; Lawrence, M. S.; Lin, P.; Meier, S.; Noble, M. S.; Saksena, G.; Voet, D.; Zhang, H.; Bernard, B.; Chambwe, N.; Dhankani, V.; Knijnenburg, T.; Kramer, R.; Leinonen, K.; Liu, Y.; Miller, M.; Reynolds, S.; Shmulevich, I.; Thorsson, V.; Zhang, W.; Akbani, R.; Broom, B. M.; Hegde, A. M.; Ju, Z.; Kanchi, R. S.; Korkut, A.; Li, J.; Liang, H.; Ling, S.; Liu, W.; Lu, Y.; Mills, G. B.; Ng, K. S.; Rao, A.; Ryan, M.; Wang, J.; Weinstein, J. N.; Zhang, J.; Abeshouse, A.; Armenia, J.; Chakravarty, D.; Chatila, W. K.; de Bruijn, I.; Gao, J.; Gross, B. E.; Heins, Z. J.; Kundra, R.; La, K.; Ladanyi, M.; Luna, A.; Nissan, M. G.; Ochoa, A.; Phillips, S. M.; Reznik, E.; Sanchez-Vega, F.; Sander, C.; Schultz, N.; Sheridan, R.; Sumer, S. O.; Sun, Y.; Taylor, B. S.; Wang, J.; Zhang, H.; Anur, P.; Peto, M.; Spellman, P.; Benz, C.; Stuart, J. M.; Wong, C. K.; Yau, C.; Hayes, D. N.; Parker, J. S.; Wilkerson, M. D.; Ally, A.; Balasundaram, M.; Bowlby, R.; Brooks, D.; Carlsen, R.; Chuah, E.; Dhalla, N.; Jones, S. J. M.; Kasaian, K.; Lee, D.; Ma, Y.; Marra, M. A.; Mayo, M.; Moore, R. A.; Mungall, A. J.; Mungall, K.; Robertson, A. G.; Sadeghi, S.; Schein, J. E.; Sipahimalani, P.; Tam, A.; Thiessen, N.; Tse, K.; Wong, T.; Berger, A. C.; Beroukhim, R.; Cherniack, A. D.; Cibulskis, C.; Gabriel, S. B.; Ha, G.; Meyerson, M.; Schumacher, S. E.; Shih, J.; Kucherlapati, M. H.; Kucherlapati, R. S.; Baylin, S.; Cope, L.; Danilova, L.; Bootwalla, M. S.; Lai, P. H.; Maglinte, D. T.; Van Den Berg, D. J.; Weisenberger, D. J.; Auman, J. T.; Balu, S.; Bodenheimer, T.; Fan, C.; Hoadley, K. A.; Hoyle, A. P.; Jefferys, S. R.; Jones, C. D.; Meng, S.; Mieczkowski, P. A.; Mose, L. E.; Perou, A. H.; Perou, C. M.; Roach, J.; Shi, Y.; Simons, J. V.; Skelly, T.; Soloway, M. G.; Tan, D.; Veluvolu, U.; Fan, H.; Hinoue, T.; Laird, P. W.; Shen, H.; Zhou, W.; Bellair, M.; Chang, K.; Covington, K.; Creighton, C. J.; Dinh, H.; Doddapaneni, H. V.; Donehower, L. A.; Drummond, J.; Gibbs, R. A.; Glenn, R.; Hale, W.; Han, Y.; Hu, J.; Korchina, V.; Lee, S.; Lewis, L.; Li, W.; Liu, X.; Morgan, M.; Morton, D.; Muzny, D.; Santibanez, J.; Sheth, M.; Shinbrot, E.; Wang, L.; Wang, M.; Wheeler, D. A.; Xi, L.; Zhao, F.; Hess, J.; Appelbaum, E. L.; Bailey, M.; Cordes, M. G.; Ding, L.; Fronick, C. C.; Fulton, L. A.; Fulton, R. S.; Kandoth, C.; Mardis, E. R.; McLellan, M. D.; Miller, C. A.; Schmidt, H. K.; Wilson, R. K.; Crain, D.; Curley, E.; Gardner, J.; Lau, K.; Mallery, D.; Morris, S.; Paulauskis, J.; Penny, R.; Shelton, C.; Shelton, T.; Sherman, M.; Thompson, E.; Yena, P.; Bowen, J.; Gastier-Foster, J. M.; Gerken, M.; Leraas, K. M.; Lichtenberg, T. M.; Ramirez, N. C.; Wise, L.; Zmuda, E.; Corcoran, N.; Costello, T.; Hovens, C.; Carvalho, A. L.; de Carvalho, A. C.; Fregnani, J. H.; Longatto-Filho, A.; Reis, R. M.; Scapulatempo-Neto, C.; Silveira, H. C. S.; Vidal, D. O.; Burnette, A.; Eschbacher, J.; Hermes, B.; Noss, A.; Singh, R.; Anderson, M. L.; Castro, P. D.; Ittmann, M.; Huntsman, D.; Kohl, B.; Le, X.; Thorp, R.; Andry, C.; Duffy, E. R.; Lyadov, V.; Paklina, O.; Setdikova, G.; Shabunin, A.; Tavobilov, M.; McPherson, C.; Warnick, R.; Berkowitz, R.; Cramer, D.; Feltmate, C.; Horowitz, N.; Kibel, A.; Muto, M.; Raut, C. P.; Malykh, A.; Barnholtz-Sloan, J. S.; Barrett, W.; Devine, K.; Fulop, J.; Ostrom, Q. T.; Shimmel, K.; Wolinsky, Y.; Sloan, A. E.; De Rose, A.; Giuliante, F.; Goodman, M.; Karlan, B. Y.; Hagedorn, C. H.; Eckman, J.; Harr, J.; Myers, J.; Tucker, K.; Zach, L. A.; Deyarmin, B.; Hu, H.; Kvecher, L.; Larson, C.; Mural, R. J.; Somiari, S.; Vicha, A.; Zelinka, T.; Bennett, J.; Iacocca, M.; Rabeno, B.; Swanson, P.; Latour, M.; Lacombe, L.; Têtu, B.; Bergeron, A.; McGraw, M.; Staugaitis, S. M.; Chabot, J.; Hibshoosh, H.; Sepulveda, A.; Su, T.; Wang, T.; Potapova, O.; Voronina, O.; Desjardins, L.; Mariani, O.; Roman-Roman, S.; Sastre, X.; Stern, M. H.; Cheng, F.; Signoretti, S.; Berchuck, A.; Bigner, D.; Lipp, E.; Marks, J.; McCall, S.; McLendon, R.; Secord, A.; Sharp, A.; Behera, M.; Brat, D. J.; Chen, A.; Delman, K.; Force, S.; Khuri, F.; Magliocca, K.; Maithel, S.; Olson, J. J.; Owonikoko, T.; Pickens, A.; Ramalingam, S.; Shin, D. M.; Sica, G.; Van Meir, E. G.; Zhang, H.; Eijckenboom, W.; Gillis, A.; Korpershoek, E.; Looijenga, L.; Oosterhuis, W.; Stoop, H.; van Kessel, K. E.; Zwarthoff, E. C.; Calatozzolo, C.; Cuppini, L.; Cuzzubbo, S.; DiMeco, F.; Finocchiaro, G.; Mattei, L.; Perin, A.; Pollo, B.; Chen, C.; Houck, J.; Lohavanichbutr, P.; Hartmann, A.; Stoehr, C.; Stoehr, R.; Taubert, H.; Wach, S.; Wullich, B.; Kycler, W.; Murawa, D.; Wiznerowicz, M.; Chung, K.; Edenfield, W. J.; Martin, J.; Baudin, E.; Bubley, G.; Bueno, R.; De Rienzo, A.; Richards, W. G.; Kalkanis, S.; Mikkelsen, T.; Noushmehr, H.; Scarpace, L.; Girard, N.; Aymerich, M.; Campo, E.; Giné, E.; Guillermo, A. L.; Van Bang, N.; Hanh, P. T.; Phu, B. D.; Tang, Y.; Colman, H.; Evason, K.; Dottino, P. R.; Martignetti, J. A.; Gabra, H.; Juhl, H.; Akeredolu, T.; Stepa, S.; Hoon, D.; Ahn, K.; Kang, K. J.; Beuschlein, F.; Breggia, A.; Birrer, M.; Bell, D.; Borad, M.; Bryce, A. H.; Castle, E.; Chandan, V.; Cheville, J.; Copland, J. A.; Farnell, M.; Flotte, T.; Giama, N.; Ho, T.; Kendrick, M.; Kocher, J. P.; Kopp, K.; Moser, C.; Nagorney, D.; O’Brien, D.; O’Neill, B. P.; Patel, T.; Petersen, G.; Que, F.; Rivera, M.; Roberts, L.; Smallridge, R.; Smyrk, T.; Stanton, M.; Thompson, R. H.; Torbenson, M.; Yang, J. D.; Zhang, L.; Brimo, F.; Ajani, J. A.; Gonzalez, A. M. A.; Behrens, C.; Bondaruk, J.; Broaddus, R.; Czerniak, B.; Esmaeli, B.; Fujimoto, J.; Gershenwald, J.; Guo, C.; Lazar, A. J.; Logothetis, C.; Meric-Bernstam, F.; Moran, C.; Ramondetta, L.; Rice, D.; Sood, A.; Tamboli, P.; Thompson, T.; Troncoso, P.; Tsao, A.; Wistuba, I.; Carter, C.; Haydu, L.; Hersey, P.; Jakrot, V.; Kakavand, H.; Kefford, R.; Lee, K.; Long, G.; Mann, G.; Quinn, M.; Saw, R.; Scolyer, R.; Shannon, K.; Spillane, A.; Stretch, onathan; Synott, M.; Thompson, J.; Wilmott, J.; Al-Ahmadie, H.; Chan, T. A.; Ghossein, R.; Gopalan, A.; Levine, D. A.; Reuter, V.; Singer, S.; Singh, B.; Tien, N. V.; Broudy, T.; Mirsaidi, C.; Nair, P.; Drwiega, P.; Miller, J.; Smith, J.; Zaren, H.; Park, J. W.; Hung, N. P.; Kebebew, E.; Linehan, W. M.; Metwalli, A. R.; Pacak, K.; Pinto, P. A.; Schiffman, M.; Schmidt, L. S.; Vocke, C. D.; Wentzensen, N.; Worrell, R.; Yang, H.; Moncrieff, M.; Goparaju, C.; Melamed, J.; Pass, H.; Botnariuc, N.; Caraman, I.; Cernat, M.; Chemencedji, I.; Clipca, A.; Doruc, S.; Gorincioi, G.; Mura, S.; Pirtac, M.; Stancul, I.; Tcaciuc, D.; Albert, M.; Alexopoulou, I.; Arnaout, A.; Bartlett, J.; Engel, J.; Gilbert, S.; Parfitt, J.; Sekhon, H.; Thomas, G.; Rassl, D. M.; Rintoul, R. C.; Bifulco, C.; Tamakawa, R.; Urba, W.; Hayward, N.; Timmers, H.; Antenucci, A.; Facciolo, F.; Grazi, G.; Marino, M.; Merola, R.; de Krijger, R.; Gimenez-Roqueplo, A. P.; Piché, A.; Chevalier, S.; McKercher, G.; Birsoy, K.; Barnett, G.; Brewer, C.; Farver, C.; Naska, T.; Pennell, N. A.; Raymond, D.; Schilero, C.; Smolenski, K.; Williams, F.; Morrison, C.; Borgia, J. A.; Liptay, M. J.; Pool, M.; Seder, C. W.; Junker, K.; Omberg, L.; Dinkin, M.; Manikhas, G.; Alvaro, D.; Bragazzi, M. C.; Cardinale, V.; Carpino, G.; Gaudio, E.; Chesla, D.; Cottingham, S.; Dubina, M.; Moiseenko, F.; Dhanasekaran, R.; Becker, K. F.; Janssen, K. P.; Slotta-Huspenina, J.; Abdel-Rahman, M. H.; Aziz, D.; Bell, S.; Cebulla, C. M.; Davis, A.; Duell, R.; Elder, J. B.; Hilty, J.; Kumar, B.; Lang, J.; Lehman, N. L.; Mandt, R.; Nguyen, P.; Pilarski, R.; Rai, K.; Schoenfield, L.; Senecal, K.; Wakely, P.; Hansen, P.; Lechan, R.; Powers, J.; Tischler, A.; Grizzle, W. E.; Sexton, K. C.; Kastl, A.; Henderson, J.; Porten, S.; Waldmann, J.; Fassnacht, M.; Asa, S. L.; Schadendorf, D.; Couce, M.; Graefen, M.; Huland, H.; Sauter, G.; Schlomm, T.; Simon, R.; Tennstedt, P.; Olabode, O.; Nelson, M.; Bathe, O.; Carroll, P. R.; Chan, J. M.; Disaia, P.; Glenn, P.; Kelley, R. K.; Landen, C. N.; Phillips, J.; Prados, M.; Simko, J.; Smith-McCune, K.; VandenBerg, S.; Roggin, K.; Fehrenbach, A.; Kendler, A.; Sifri, S.; Steele, R.; Jimeno, A.; Carey, F.; Forgie, I.; Mannelli, M.; Carney, M.; Hernandez, B.; Campos, B.; Herold-Mende, C.; Jungk, C.; Unterberg, A.; von Deimling, A.; Bossler, A.; Galbraith, J.; Jacobus, L.; Knudson, M.; Knutson, T.; Ma, D.; Milhem, M.; Sigmund, R.; Godwin, A. K.; Madan, R.; Rosenthal, H. G.; Adebamowo, C.; Adebamowo, S. N.; Boussioutas, A.; Beer, D.; Giordano, T.; Mes-Masson, A. M.; Saad, F.; Bocklage, T.; Landrum, L.; Mannel, R.; Moore, K.; Moxley, K.; Postier, R.; Walker, J.; Zuna, R.; Feldman, M.; Valdivieso, F.; Dhir, R.; Luketich, J.; Pinero, E. M. M.; Quintero-Aguilo, M.; Carlotti, C. G.; Dos Santos, J. S.; Kemp, R.; Sankarankuty, A.; Tirapelli, D.; Catto, J.; Agnew, K.; Swisher, E.; Creaney, J.; Robinson, B.; Shelley, C. S.; Godwin, E. M.; Kendall, S.; Shipman, C.; Bradford, C.; Carey, T.; Haddad, A.; Moyer, J.; Peterson, L.; Prince, M.; Rozek, L.; Wolf, G.; Bowman, R.; Fong, K. M.; Yang, I.; Korst, R.; Rathmell, W. K.; Fantacone-Campbell, J. L.; Hooke, J. A.; Kovatich, A. J.; Shriver, C. D.; DiPersio, J.; Drake, B.; Govindan, R.; Heath, S.; Ley, T.; Van Tine, B.; Westervelt, P.; Rubin, M. A.; Lee, J. Il; Aredes, N. D.; Mariamidze, A.; Serody, J. S.; Demicco, E. G.; Disis, M. L.; Vincent, B. G.; Shmulevich, llya. The Immune Landscape of Cancer. Immunity 2018, 48 (4), 812–830.e14. 10.1016/j.immuni.2018.03.023.

(3) Anderson, N. M.; Simon, M. C. The Tumor Microenvironment. Curr. Biol. CB 2020, 30 (16), R921–R925. 10.1016/j.cub.2020.06.081.

(4) Kusmartsev, S.; Su, Z.; Heiser, A.; Dannull, J.; Eruslanov, E.; Kübler, H.; Yancey, D.; Dahm, P.; Vieweg, J. Reversal of Myeloid Cell - Mediated Immunosuppression in Patients with Metastatic Renal Cell Carcinoma. Clin. Cancer Res. 2008, 14 (24), 8270–8278. 10.1158/1078-0432.CCR-08-0165.

(5) Gabrilovich, D. I.; Ostrand-Rosenberg, S.; Bronte, V. Coordinated Regulation of Myeloid Cells by Tumours. Nat. Rev. Immunol. 2012, 12 (4), 253–268. 10.1038/nri3175.

(6) Kuan, E. L.; Ziegler, S. F. A Tumor-Myeloid Cell Axis, Mediated via the Cytokines IL-1α and TSLP, Promotes the Progression of Breast Cancer. Nat. Immunol. 2018, 19 (4), 366–374. 10.1038/s41590-018-0066-6.

(7) Morello, S.; Pinto, A.; Blandizzi, C.; Antonioli, L. Myeloid Cells in the Tumor Microenvironment: Role of Adenosine. OncoImmunology 2016, 5 (3), 1–8. 10.1080/2162402X.2015.1108515.

(8) Ushach, I.; Zlotnik, A. Biological Role of Granulocyte Macrophage Colony-Stimulating Factor (GM-CSF) and Macrophage Colony-Stimulating Factor (M-CSF) on Cells of the Myeloid Lineage. J. Leukoc. Biol. 2016, 100 (3), 481–489. 10.1189/JLB.3RU0316-144R.

(9) Nakatsumi, H.; Matsumoto, M.; Nakayama, K. I. Noncanonical Pathway for Regulation of CCL2 Expression by an mTORC1-FOXK1 Axis Promotes Recruitment of Tumor-Associated Macrophages. Cell Rep. 2017, 21 (9), 2471–2486. 10.1016/j.celrep.2017.11.014.

(10) Ugel, S.; De Sanctis, F.; Mandruzzato, S.; Bronte, V. Tumor-Induced Myeloid Deviation: When Myeloid-Derived Suppressor Cells Meet Tumor-Associated Macrophages. J. Clin. Invest. 2015, 125 (9), 3365–3376. 10.1172/JCI80006.

(11) Liew, P. X.; Kubes, P. The Neutrophil’s Role during Health and Disease. Physiol. Rev. 2019, 99 (2), 1223–1248. 10.1152/physrev.00012.2018.

(12) Patel, S.; Fu, S.; Mastio, J.; Dominguez, G. A.; Purohit, A.; Kossenkov, A.; Lin, C.; Alicea-Torres, K.; Sehgal, M.; Nefedova, Y.; Zhou, J.; Languino, L. R.; Clendenin, C.; Vonderheide, R. H.; Mulligan, C.; Nam, B.; Hockstein, N.; Masters, G.; Guarino, M.; Schug, Z. T.; Altieri, D. C.; Gabrilovich, D. I. Unique Pattern of Neutrophil Migration and Function during Tumor Progression. Nat. Immunol. 2018, 19 (11), 1236–1247. 10.1038/s41590-018-0229-5.

(13) Nishida, J.; Momoi, Y.; Miyakuni, K.; Tamura, Y.; Takahashi, K.; Koinuma, D.; Miyazono, K.; Ehata, S. Epigenetic Remodelling Shapes Inflammatory Renal Cancer and Neutrophil-Dependent Metastasis. Nat. Cell Biol. 2020, 22 (4), 465–475. 10.1038/s41556-020-0491-2.

(14) IS, K.; Y, G.; T, W.; H, W.; J, L.; M, J.; K, S.; Y, N.; A, G.; N, Z.; I, B.; HC, L.; MJ, T.; T, N.; W, B.; W, J.; J, A.; F, G.; J, H.; D, J.; K, W.; Y, L.; Q, M.; TF, W.; C, Z.; A, R.; A, S.; JM, R.; XH, Z. Immuno-Subtyping of Breast Cancer Reveals Distinct Myeloid Cell Profiles and Immunotherapy Resistance Mechanisms. Nat. Cell Biol. 2019, 21 (9), 1113–1126. 10.1038/S41556-019-0373-7.

(15) Liu, H.; Wang, Z.; Zhou, Y.; Yang, Y. MDSCs in Breast Cancer: An Important Enabler of Tumor Progression and an Emerging Therapeutic Target. Front. Immunol. 2023, 14 (July), 1–17. 10.3389/fimmu.2023.1199273.

(16) Zhang, Y.; Murphy, S.; Lu, X. Cancer-Cell-Intrinsic Mechanisms Regulate MDSCs through Cytokine Networks; 2023; pp 1–31. 10.1016/bs.ircmb.2022.09.001.

(17) Chesney, J. A.; Mitchell, R. A.; Yaddanapudi, K. Myeloid-Derived Suppressor Cells—a New Therapeutic Target to Overcome Resistance to Cancer Immunotherapy. J. Leukoc. Biol. 2017, 102 (3), 727–740. 10.1189/jlb.5vmr1116-458rrr.

(18) Siegel, R. L.; Giaquinto, A. N.; Jemal, A. Cancer Statistics, 2024. CA. Cancer J. Clin. 2024, 74 (1), 12–49. 10.3322/CAAC.21820.

(19) Wang, S.; Gao, J.; Lei, Q.; Rozengurt, N.; Pritchard, C.; Jiao, J.; Thomas, G. V.; Li, G.; Roy-Burman, P.; Nelson, P. S.; Liu, X.; Wu, H. Prostate-Specific Deletion of the Murine Pten Tumor Suppressor Gene Leads to Metastatic Prostate Cancer. Cancer Cell 2003, 4 (3), 209–221. 10.1016/S1535-6108(03)00215-0.

(20) Chen, Z.; Trotman, L. C.; Shaffer, D.; Lin, H.-K.; Dotan, Z. A.; Niki, M.; Koutcher, J. A.; Scher, H. I.; Ludwig, T.; Gerald, W.; Cordon-Cardo, C.; Paolo Pandolfi, P. Crucial Role of P53-Dependent Cellular Senescence in Suppression of Pten-Deficient Tumorigenesis. Nature 2005, 436 (7051), 725–730. 10.1038/nature03918.

(21) Ding, Z.; Wu, C. J.; Chu, G. C.; Xiao, Y.; Ho, D.; Zhang, J.; Perry, S. R.; Labrot, E. S.; Wu, X.; Lis, R.; Hoshida, Y.; Hiller, D.; Hu, B.; Jiang, S.; Zheng, H.; Stegh, A. H.; Scott, K. L.; Signoretti, S.; Bardeesy, N.; Wang, Y. A.; Hill, D. E.; Golub, T. R.; Stampfer, M. J.; Wong, W. H.; Loda, M.; Mucci, L.; Chin, L.; Depinho, R. A. SMAD4-Dependent Barrier Constrains Prostate Cancer Growth and Metastatic Progression. Nature 2011, 470 (7333), 269–276. 10.1038/nature09677.

(22) Zeng, L.; Rowland, R. G.; Lele, S. M.; Kyprianou, N. Apoptosis Incidence and Protein Expression of P53, TGF-β Receptor II, p27Kip1, and Smad4 in Benign, Premalignant, and Malignant Human Prostate. Hum. Pathol. 2004, 35 (3), 290–297. 10.1016/j.humpath.2003.11.001.

(23) Wang, G.; Lu, X.; Dey, P.; Deng, P.; Wu, C. C.; Jiang, S.; Fang, Z.; Zhao, K.; Konaparthi, R.; Hua, S.; Zhang, J.; Li-Ning-Tapia, E. M.; Kapoor, A.; Wu, C.-J.; Patel, N. B.; Guo, Z.; Ramamoorthy, V.; Tieu, T. N.; Heffernan, T.; Zhao, D.; Shang, X.; Khadka, S.; Hou, P.; Hu, B.; Jin, E.-J.; Yao, W.; Pan, X.; Ding, Z.; Shi, Y.; Li, L.; Chang, Q.; Troncoso, P.; Logothetis, C. J.; McArthur, M. J.; Chin, L.; Wang, Y. A.; DePinho, R. A. Targeting YAP-Dependent MDSC Infiltration Impairs Tumor Progression. Cancer Discov. 2016, 6 (1), 80–95. 10.1158/2159-8290.CD-15-0224.

(24) Lu, X.; Horner, J. W.; Paul, E.; Shang, X.; Troncoso, P.; Deng, P.; Jiang, S.; Chang, Q.; Spring, D. J.; Sharma, P.; Zebala, J. A.; Maeda, D. Y.; Wang, Y. A.; Depinho, R. A. Effective Combinatorial Immunotherapy for Castration-Resistant Prostate Cancer. Nature 2017, 543 (7647), 728–732. 10.1038/nature21676.

(25) Srivastava, A. K.; Guadagnin, G.; Cappello, P.; Novelli, F. Post-Translational Modifications in Tumor-Associated Antigens as a Platform for Novel Immuno-Oncology Therapies. Cancers 2022, 15 (1), 138. 10.3390/CANCERS15010138.

(26) Audia, J. E.; Campbell, R. M. Histone Modifications and Cancer. Cold Spring Harb. Perspect. Biol. 2016, 8 (4), a019521. 10.1101/CSHPERSPECT.A019521.

(27) Dutta, H.; Jain, N. Post-Translational Modifications and Their Implications in Cancer. Front. Oncol. 2023, 13, 1240115. 10.3389/FONC.2023.1240115.

(28) Yuzhalin, A. E. Citrullination in Cancer. Cancer Research. American Association for Cancer Research Inc. 2019, pp 1274–1284. 10.1158/0008-5472.CAN-18-2797.

(29) Zhu, D.; Zhang, Y.; Wang, S. Histone Citrullination: A New Target for Tumors. Mol. Cancer 2021, 20 (1), 1–17. 10.1186/S12943-021-01373-Z/TABLES/2.

(30) Sharma, P.; Azebi, S.; England, P.; Christensen, T.; Møller-Larsen, A.; Petersen, T.; Batsché, E.; Muchardt, C. Citrullination of Histone H3 Interferes with HP1-Mediated Transcriptional Repression. PLoS Genet. 2012, 8 (9). 10.1371/JOURNAL.PGEN.1002934.

(31) Sun, B.; Dwivedi, N.; Bechtel, T. J.; Paulsen, J. L.; Muth, A.; Bawadekar, M.; Li, G.; Thompson, P. R.; Shelef, M. A.; Schiffer, C. A.; Weerapana, E.; Ho, I.-C. Citrullination of NF-κB P65 Promotes Its Nuclear Localization and TLR-Induced Expression of IL-1κ and TNFκ; 2017; Vol. 2.

(32) Cherrington, B. D.; Zhang, X.; McElwee, J. L.; Morency, E.; Anguish, L. J.; Coonrod, S. A. Potential Role for PAD2 in Gene Regulation in Breast Cancer Cells. PloS One 2012, 7 (7). 10.1371/JOURNAL.PONE.0041242.

(33) McElwee, J. L.; Mohanan, S.; Griffith, O. L.; Breuer, H. C.; Anguish, L. J.; Cherrington, B. D.; Palmer, A. M.; Howe, L. R.; Subramanian, V.; Causey, C. P.; Thompson, P. R.; Gray, J. W.; Coonrod, S. A. Identification of PADI2 as a Potential Breast Cancer Biomarker and Therapeutic Target. BMC Cancer 2012, 12. 10.1186/1471-2407-12-500.

(34) Xue, T.; Liu, X.; Zhang, M.; Qiukai, E.; Liu, S.; Zou, M.; Li, Y.; Ma, Z.; Han, Y.; Thompson, P.; Zhang, X. PADI2-Catalyzed MEK1 Citrullination Activates ERK1/2 and Promotes IGF2BP1-Mediated SOX2 mRNA Stability in Endometrial Cancer. Adv. Sci. 2021, 8 (6), 2002831. 10.1002/ADVS.202002831.

(35) Wang, L.; Song, G.; Zhang, X.; Feng, T.; Pan, J.; Chen, W.; Yang, M.; Bai, X.; Pang, Y.; Yu, J.; Han, J.; Han, B. PADI2-Mediated Citrullination Promotes Prostate Cancer Progression. Cancer Res. 2017, 77 (21), 5755–5768. 10.1158/0008-5472.CAN-17-0150.

(36) Zhu, Y.; Zhao, Y.; Wen, J.; Liu, S.; Huang, T.; Hatial, I.; Peng, X.; Al Janabi, H.; Huang, G.; Mittlesteadt, J.; Cheng, M.; Bhardwaj, A.; Ashfeld, B. L.; Kao, K. R.; Maeda, D. Y.; Dai, X.; Wiest, O.; Blagg, B. S. J.; Lu, X.; Cheng, L.; Wan, J.; Lu, X. Targeting the Chromatin Effector Pygo2 Promotes Cytotoxic T Cell Responses and Overcomes Immunotherapy Resistance in Prostate Cancer. Sci. Immunol. 2023, 8 (81), eade4656. 10.1126/sciimmunol.ade4656.

(37) Robinson, M. D.; McCarthy, D. J.; Smyth, G. K. edgeR: A Bioconductor Package for Differential Expression Analysis of Digital Gene Expression Data. Bioinforma. Oxf. Engl. 2010, 26 (1), 139–140. 10.1093/bioinformatics/btp616.

(38) Jin, S.; Guerrero-Juarez, C. F.; Zhang, L.; Chang, I.; Ramos, R.; Kuan, C.-H.; Myung, P.; Plikus, M. V.; Nie, Q. Inference and Analysis of Cell-Cell Communication Using CellChat. Nat. Commun. 2021, 12 (1), 1088. 10.1038/s41467-021-21246-9.

(39) Langmead, B.; Salzberg, S. L. Fast Gapped-Read Alignment with Bowtie 2. Nat. Methods 2012, 9 (4), 357–359. 10.1038/nmeth.1923.

(40) Li, H.; Handsaker, B.; Wysoker, A.; Fennell, T.; Ruan, J.; Homer, N.; Marth, G.; Abecasis, G.; Durbin, R.; Subgroup, 1000 Genome Project Data Processing. The Sequence Alignment/Map Format and SAMtools. Bioinforma. Oxf. Engl. 2009, 25 (16), 2078–2079. 10.1093/bioinformatics/btp352.

(41) Zhang, Y.; Liu, T.; Meyer, C. A.; Eeckhoute, J.; Johnson, D. S.; Bernstein, B. E.; Nusbaum, C.; Myers, R. M.; Brown, M.; Li, W.; Liu, X. S. Model-Based Analysis of ChIP-Seq (MACS). Genome Biol. 2008, 9 (9), R137. 10.1186/gb-2008-9-9-r137.

(42) Quinlan, A. R.; Hall, I. M. BEDTools: A Flexible Suite of Utilities for Comparing Genomic Features. Bioinforma. Oxf. Engl. 2010, 26 (6), 841–842. 10.1093/bioinformatics/btq033.

(43) Yu, G.; Wang, L.-G.; He, Q.-Y. ChIPseeker: An R/Bioconductor Package for ChIP Peak Annotation, Comparison and Visualization. Bioinforma. Oxf. Engl. 2015, 31 (14), 2382–2383. 10.1093/bioinformatics/btv145.

(44) Risso, D.; Schwartz, K.; Sherlock, G.; Dudoit, S. GC-Content Normalization for RNA-Seq Data. BMC Bioinformatics 2011, 12, 480. 10.1186/1471-2105-12-480.

(45) IS, K.; Y, G.; T, W.; H, W.; J, L.; M, J.; K, S.; Y, N.; A, G.; N, Z.; I, B.; HC, L.; MJ, T.; T, N.; W, B.; W, J.; J, A.; F, G.; J, H.; D, J.; K, W.; Y, L.; Q, M.; TF, W.; C, Z.; A, R.; A, S.; JM, R.; XH, Z. Immuno-Subtyping of Breast Cancer Reveals Distinct Myeloid Cell Profiles and Immunotherapy Resistance Mechanisms. Nat. Cell Biol. 2019, 21 (9), 1113–1126. 10.1038/S41556-019-0373-7.

(46) Wang, Y.; Wysocka, J.; Sayegh, J.; Lee, Y. H.; Pertin, J. R.; Leonelli, L.; Sonbuchner, L. S.; McDonald, C. H.; Cook, R. G.; Dou, Y.; Roeder, R. G.; Clarke, S.; Stallcup, M. R.; Allis, C. D.; Coonrod, S. A. Human PAD4 Regulates Histone Arginine Methylation Levels via Demethylimination. Science 2004, 306 (5694), 279–283. 10.1126/SCIENCE.1101400.

(47) Beato, M.; Sharma, P. Peptidyl Arginine Deiminase 2 (PADI2)-Mediated Arginine Citrullination Modulates Transcription in Cancer. Int. J. Mol. Sci. 2020, 21 (4). 10.3390/IJMS21041351.

(48) Zhang, X.; Bolt, M.; Guertin, M. J.; Chen, W.; Zhang, S.; Cherrington, B. D.; Slade, D. J.; Dreyton, C. J.; Subramanian, V.; Bicker, K. L.; Thompson, P. R.; Mancini, M. A.; Lis, J. T.; Coonrod, S. A. Peptidylarginine Deiminase 2-Catalyzed Histone H3 Arginine 26 Citrullination Facilitates Estrogen Receptor α Target Gene Activation. Proc. Natl. Acad. Sci. U. S. A. 2012, 109 (33), 13331–13336. 10.1073/pnas.1203280109.

(49) Taylor, B. S.; Schultz, N.; Hieronymus, H.; Gopalan, A.; Xiao, Y.; Carver, B. S.; Arora, V. K.; Kaushik, P.; Cerami, E.; Reva, B.; Antipin, Y.; Mitsiades, N.; Landers, T.; Dolgalev, I.; Major, J. E.; Wilson, M.; Socci, N. D.; Lash, A. E.; Heguy, A.; Eastham, J. A.; Scher, H. I.; Reuter, V. E.; Scardino, P. T.; Sander, C.; Sawyers, C. L.; Gerald, W. L. Integrative Genomic Profiling of Human Prostate Cancer. Cancer Cell 2010, 18 (1), 11–22. 10.1016/j.ccr.2010.05.026.

(50) Condamine, T.; Dominguez, G. A.; Youn, J. I.; Kossenkov, A. V.; Mony, S.; Alicea-Torres, K.; Tcyganov, E.; Hashimoto, A.; Nefedova, Y.; Lin, C.; Partlova, S.; Garfall, A.; Vogl, D. T.; Xu, X.; Knight, S. C.; Malietzis, G.; Lee, G. H.; Eruslanov, E.; Albelda, S. M.; Wang, X.; Mehta, J. L.; Bewtra, M.; Rustgi, A.; Hockstein, N.; Witt, R.; Masters, G.; Nam, B.; Smirnov, D.; Sepulveda, M. A.; Gabrilovich, D. I. Lectin-Type Oxidized LDL Receptor-1 Distinguishes Population of Human Polymorphonuclear Myeloid-Derived Suppressor Cells in Cancer Patients. Sci. Immunol. 2016, 1 (2), aaf8943–aaf8943. 10.1126/sciimmunol.aaf8943.

