## Supplemental Figures S1-S7 for "An oncogenotype-immunophenotype paradigm governing the myeloid landscape in genetically engineered mouse models of prostate cancer"

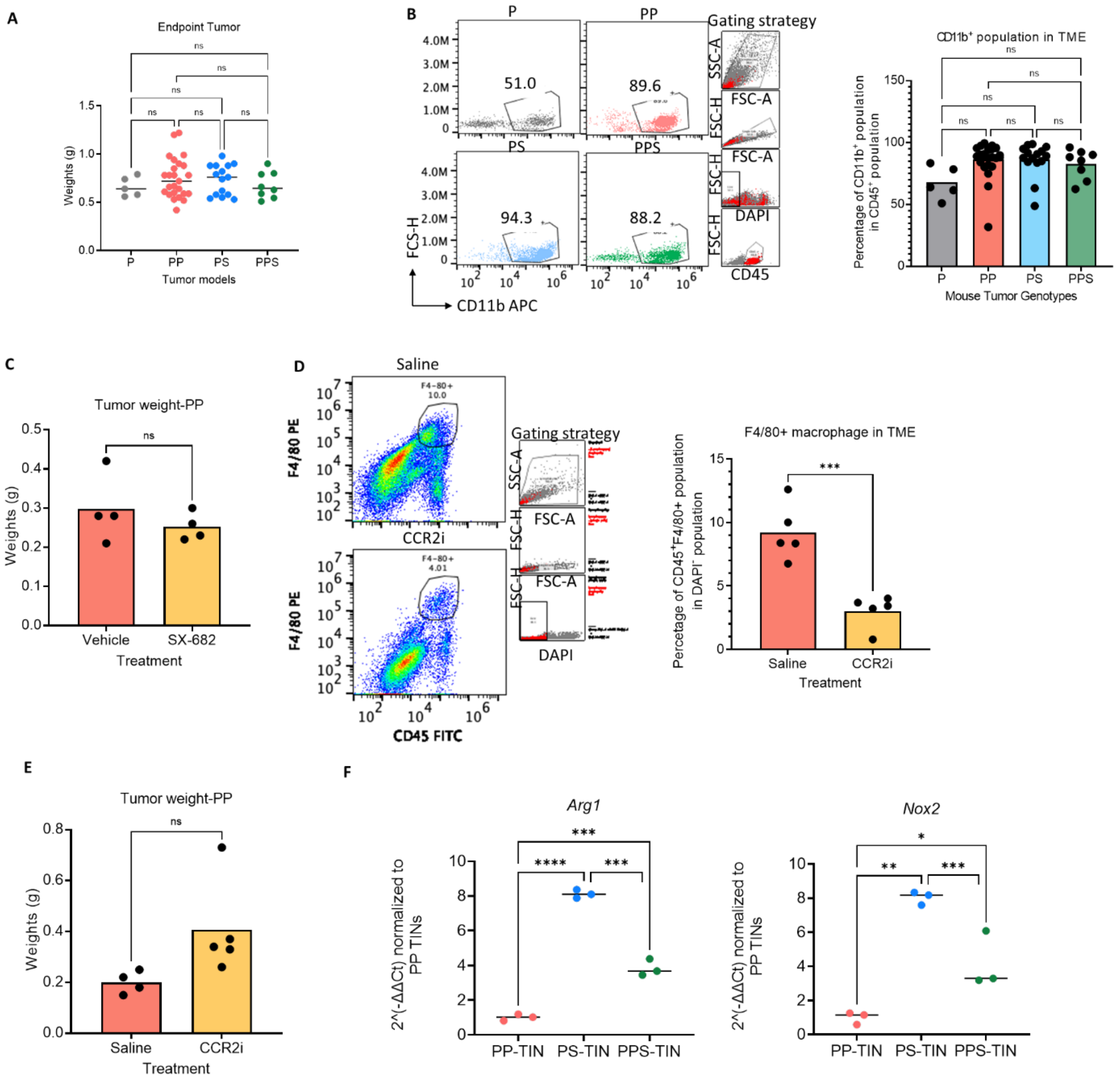


Figure S1. A distinct myeloid cell distribution across different genotypes of tumor models. (A) Illustration for the harvest ages across four prostate tumor models. Size-matched tumors (around 1 gram) were used for immune profiling. (B) FACS analyses showing infiltration $CD11b^+$ cells (myeloid cells) in four representative tumor models. Plots are gated on $CD45^+$ cells. (C) The bar plot of PP tumor weights of vehicle- and SX-682-treated tumor bearing mice. (D) FACS analyses showing TIM percentage in the tumor microenvironment of vehicle- and CCR2i-treated tumor bearing mice. Plots are gated on whole $DAPI^-$ live cells. (E) The bar plot of PP tumor weights of vehicle- and CCR2i-treated tumor bearing mice. (F) qRT-PCR results of *Arg1* and *Nox2* gene expression in tumor-associated neutrophils purified from PP, PS, PPS tumors. *ns* $p > 0.05$, * $p < 0.05$, ** $p < 0.01$,*** $p < 0.001$, **** $p < 0.0001$, Unpaired nonparametric Mann Whitney test.

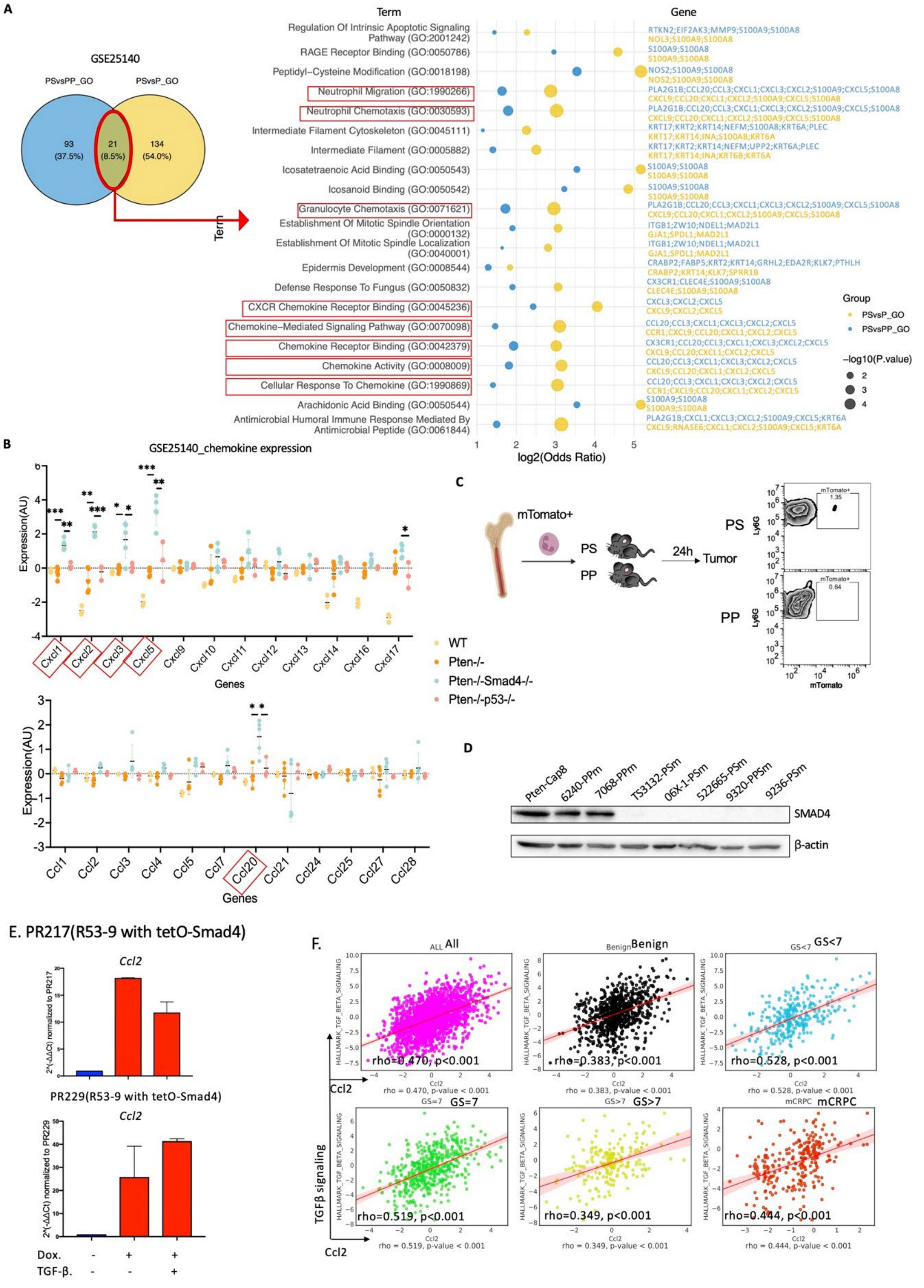


Figure S2. Tumor-intrinsic factors contribute to immunosuppression in TME. (A) Venn plot of the overlapped signaling enrichment in PS vs PP and PS vs P in GSE25140 dataset. (B) Dot plot showing the normalized value of each chemokine expression. (C) Flow cytometry analysis of mTomato$^{+}$ bone marrow neutrophil transplantation into PP- and PS-tumor bearing mice. (D) Western blotting of SMAD4 expression in the tumor-derived cell lines. (E) qRT-PCR analysis of *Ccl2* expression in PR217 and PR229 cell lines (two PS cell lines) with TGFβ signaling active or inactive. Doxycycline (Dox) was used to induce *Smad4* expression. (F) Correlation between *Ccl2* expression and the HALLMARK_TGF_BETA_SIGNALING signature in the Prostate Cancer Transcriptome Atlas (PCTA) dataset, assessed using the PCTA web tool. *ns* $p > 0.05$, * $p < 0.05$, ** $p < 0.01$, *** $p < 0.001$, **** $p < 0.0001$, Unpaired nonparametric Mann Whitney test.

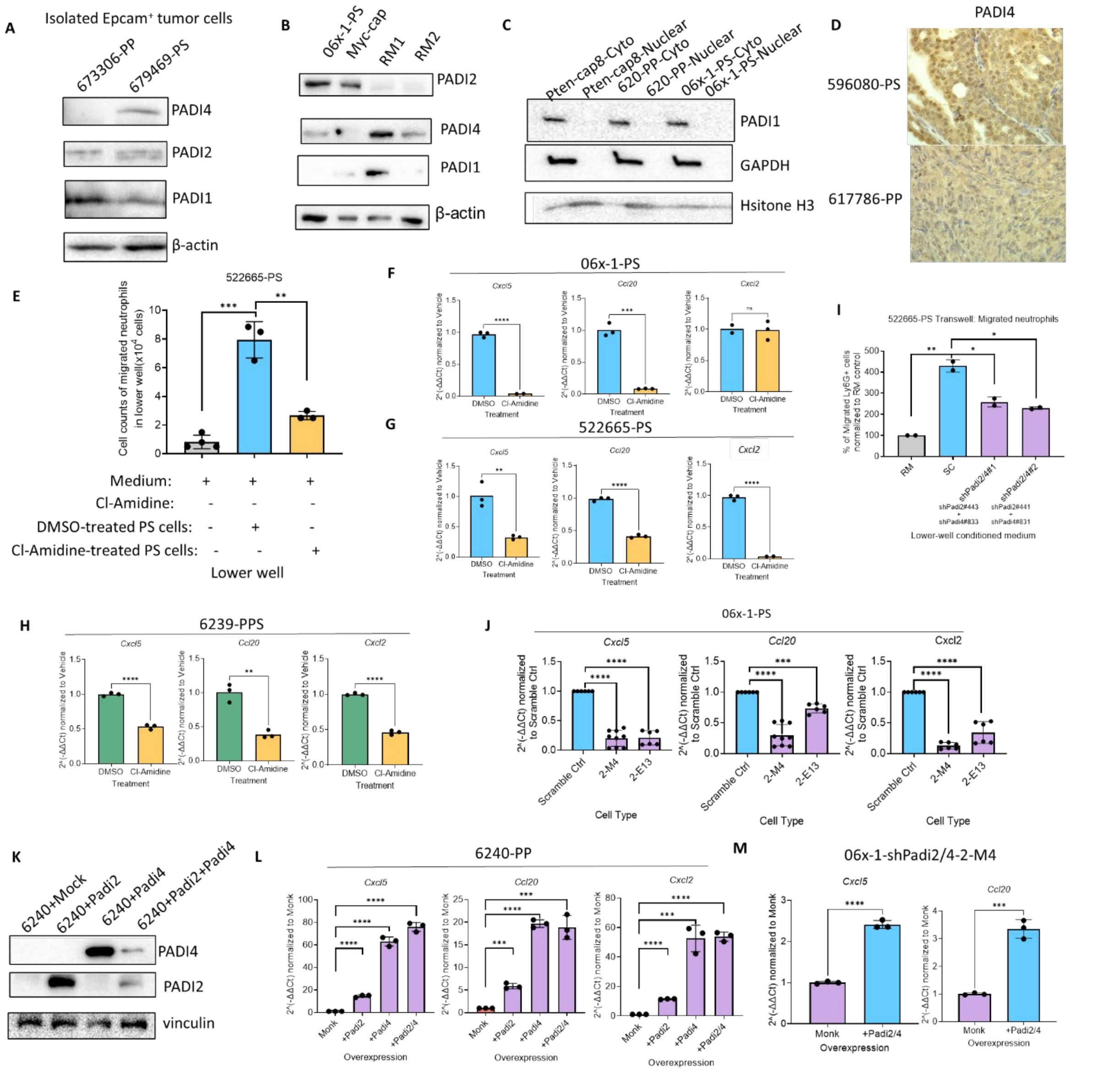

Figure S3. Protein-arginine deiminase (PADI) activity upregulated in PS and PPS tumors and its role in immunosuppressive neutrophil accumulation. (A) Western blotting of PADI1,2,4 expression in Epcam+ purified PP, PS tumor cells from tumor tissues. (B) Western blotting of PADI1, 2, 4 expressions in some other mouse prostate cancer cell lines. (C) Cell Fragmentation western blotting of PADI1 expression in Pten-cap8, 6240-PP and 06x-1-PS cell line. (D) IHC of PADI4 in PS and PP tumor tissue. (E) Transwell migration assay of migrated neutrophil count in vehicle- or Cl-Amidine- treated 522665-PS cells. (F-H) qPCR analysis of related chemokine expression (C*xcl5, Ccl20, Cxcl2*) in vehicle- or Cl-Amidine-treated 06x-1-PS(F), 522665-PS(G) and 6239-PPS(H) cells. (I) Transwell assay of migrated neutrophil count in scramble control or sh*Padi2/4*-KD of 522665-PS. (J) qPCR analysis of related chemokine expression *Cxcl5, Ccl20, Cxcl2* in scramble control or sh*Padi2/4* KD of 06x-1-PS cells. (K) Western blotting of PADI4 and PADI2 expression in overexpression of *Padi2/4* in 6240-PP cells. (L) qPCR results of chemokine expression *Cxcl5, Ccl20, Cxcl2* in overexpression of PADI2/4 in 6240-PP cells. (M) qPCR results of chemokine expression *Cxcl5, Ccl20, Cxcl2* in overexpression of *Padi2/4* in sh*Padi2/4*-2-M4 cells. *ns* $p > 0.05$, * $p < 0.05$, ** $p < 0.01$,*** $p < 0.001$, **** $p < 0.0001$, Unpaired nonparametric Mann Whitney test.

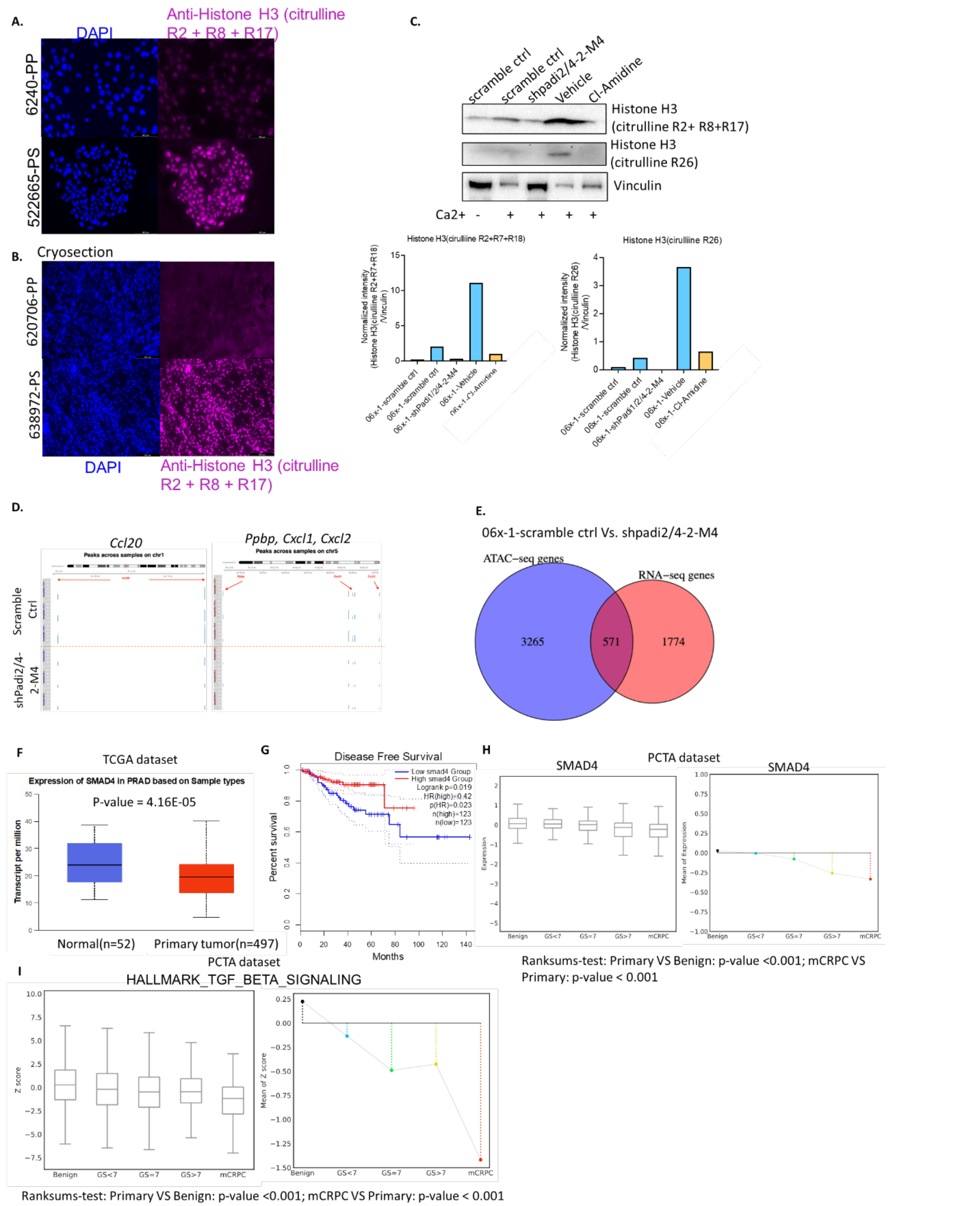


Figure S4. Enhanced histone citrullination promotes cytokine expression, leading to the recruitment of immunosuppressive neutrophils. (A-B) Immunocytochemistry analysis and quantification of Histone H3 (citrulline R2+R8+R17) expression and localization in 6240-PP and 522665-PS cells (A), and in 520706-PP and 638972-PS tumor cryosection tissues (B). (C) Western blot analysis and quantification of Histone H3 (citrulline R2+R8+R17) and Histone H3 (citrulline R26) in scramble control, sh*Padi2/*4 KD, and vehicle- or Cl-Amidine-treated 06x-1-PS cells, with or without $Ca^{2+}$. Vinculin served as a loading control. (D) ATAC-seq peaks for *Ccl20*, *Ppbp*, *Cxcl1*, and *Cxcl2* in scramble control versus sh*Padi2/4* KD 06x-1-PS cells. (E) Venn diagram displaying overlapping genes between those upregulated in RNA-seq and genes associated with open-chromatin regions from ATAC-seq in scramble control compared to sh*Padi2/4* KD 06x-1-PS cells.

(F) Box plot of SMAD4 expression levels between human normal tissue and primary tumor tissue in Prostate adenocarcinoma (PRAD) in TCGA dataset. Plot made by ULCAN. Ranksums-test between subsets (mCRPC VS Primary), P-value = <0.001; Ranksums-test between subsets(Primary VS Benign): P-value = <0.001. (G) Survival analysis based on the SMAD4 expression is depicted in a Kaplan-Meier plot via GEPIA2, covering all surveyed cancer types. The patient cohort is stratified into quartiles for this examination. (H) Box plot of SMAD4 expression(left) and mean of expression(right) among different gleason scores of prostate tumor tissue in Prostate Cancer Transcriptome Atlas(PCTA) datasets. Ranksums-test between subsets(mCRPC VS Primary), P-value = <0.001; Ranksums-test between subsets(Primary VS Benign): P-value = <0.001. (I)Box plot of Hallmark_TGF_beta_signaling enrichment(left) and mean of expression(right) among different gleason scores of prostate tumor tissue in Prostate Cancer Transcriptomics Atlas(PCTA) datasets. Ranksums-test between subsets (mCRPC VS Primary): Fold change = -1.051, P-value = <0.001; Ranksums-test between subsets(Primary VS Benign): Fold change = -0.593, P-value = <0.001.

Figure S5

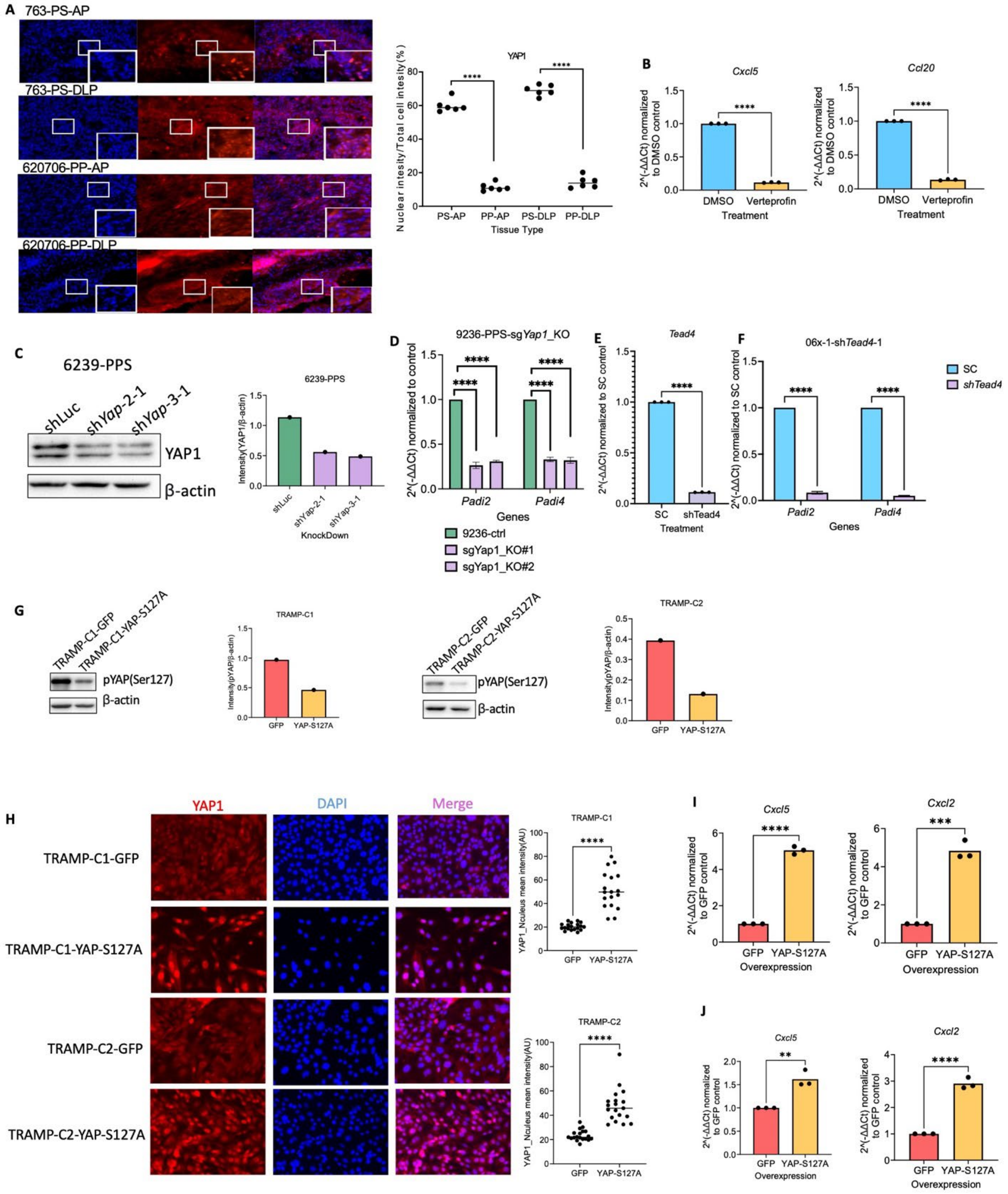


Figure S5. Smad4 ablation upregulated PADI2/4 expression via YAP signaling. (A) Immunofluorescence analysis of YAP1 in PS tumor and PP tumor tissue. (B) qPCR analysis of *Cxcl5* and *Ccl20* expression in verteporfin-treated 06x-1-PS cells and DMSO-treated 06x-1-PS cells. (C) Western Blot analysis of YAP1 expression in shLuc- and shYap1-KD 6239-PPS cells. β-actin is the loading control. (B) qPCR analysis of *Padi2* and *Padi4* expression in shScramble control- and sg*Yap1*-KO- 9236-PPS cells. (E-F) qPCR analysis of *Tead4* (E) and *Padi2* and *Padi4* (F) expression in shScramble control- and sg*Tead4*-KD- 06x-1-PS cells. (G) Western Blot analysis of phosphorylated YAP1 (pYAP(Ser127) in GFP- and YAP-S127A-TRAMP-C1 and TRAMP-C2 cells. (H) Immunocytochemistry analysis of YAP1 expression and location in GFP- and YAP-S127A-TRAMP-C1 and TRAMP-C2 cells. (I) qPCR analysis of *Cxcl5* and *Cxcl2* expression in GFP- and YAP-S127A-TRAMP-C1 and TRAMP-C2 cells. *ns* $p > 0.05$, * $p < 0.05$, ** $p < 0.01$,*** $p < 0.001$, **** $p < 0.0001$, Unpaired nonparametric Mann Whitney test.

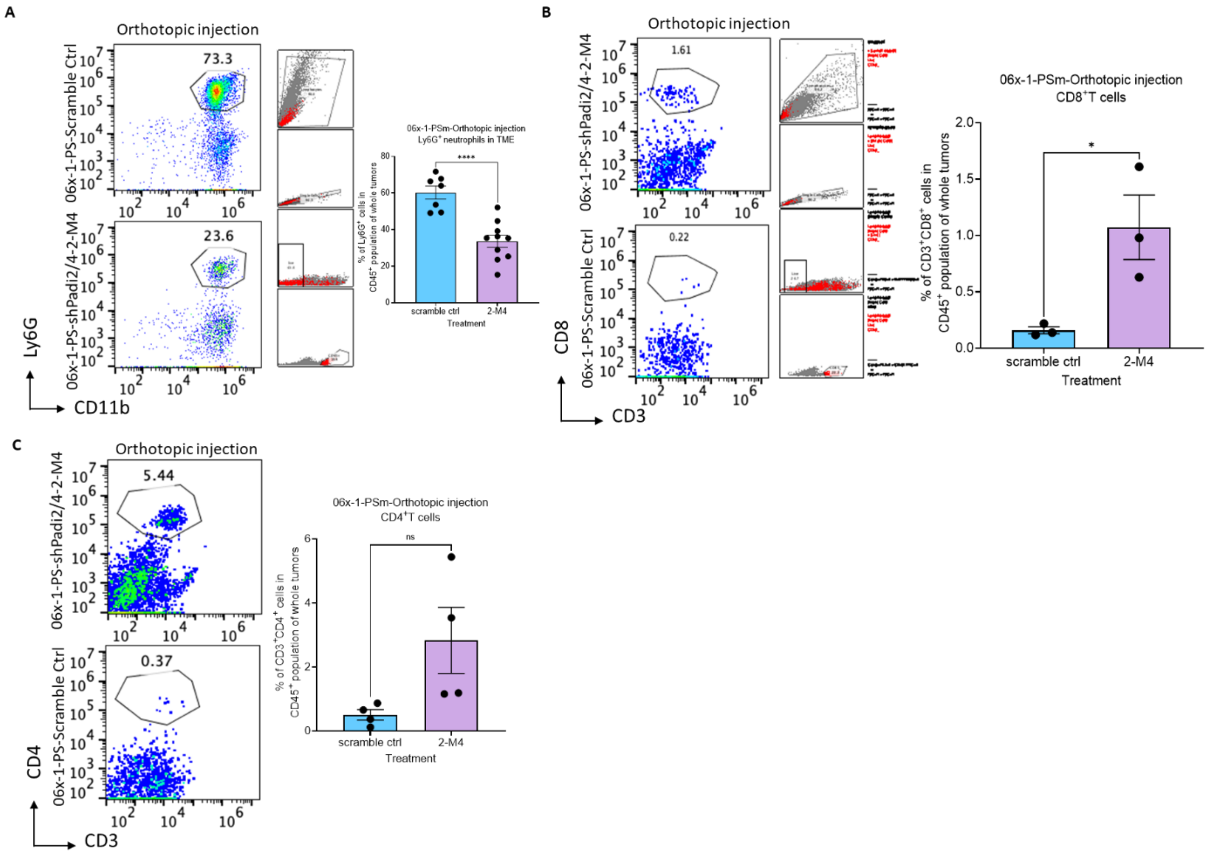


Figure S6: PADI knockdown suppresses neutrophil accumulation in prostate tumors.
(A) FACS analysis and quantification of CD11b$^+$ Ly6G$^+$ tumor-infiltrating neutrophil (TIN) percentages in the tumor microenvironment of shScramble control and sh*Padi2/4* KD prostate orthotopic 06x-1-PS tumors in C57BL/6 mice.
(B)FACS analysis and quantification of CD3$^+$ CD8$^+$ T cell percentages in the tumor microenvironment of shScramble control and sh*Padi2/4* KD prostate orthotopic 06x-1-PS tumors in C57BL/6 mice.
(C)FACS analysis and quantification of CD3$^+$ CD4$^+$ T cell percentages in the tumor microenvironment of shScramble control and sh*Padi2/4* KD prostate orthotopic 06x-1-PS tumors in C57BL/6 mice.
Statistical significance: ns $p > 0.05$, * $p < 0.05$, ** $p < 0.01$, *** $p < 0.001$, **** $p < 0.0001$, Unpaired nonparametric Mann Whitney test.

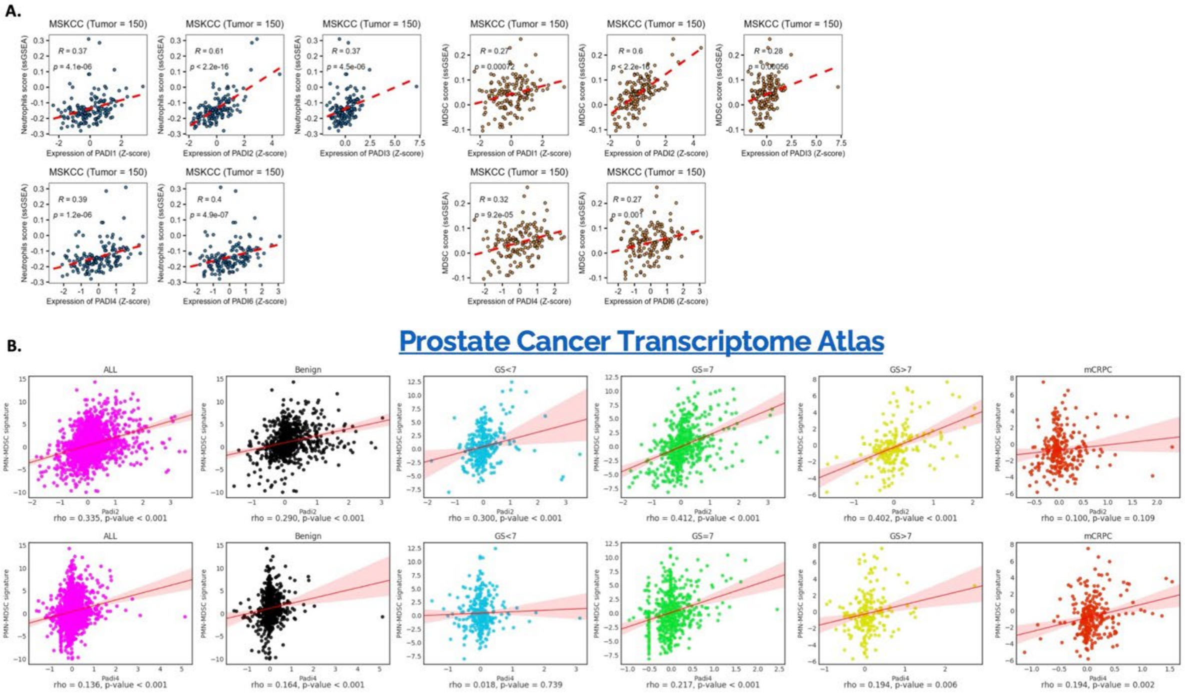


Figure S7. *PADI2/4* Is Upregulated in Human Prostate Cancer and Correlates with PMN-MDSC Recruitment. (A) Correlation analysis between neutrophil score (ssGSEA) and MDSC score (ssGSEA) with the expression of *PADI1, PADI2, PADI3*, and *PADI4* in the MSKCC Prostate Cancer Genomics Data Portal (Tumor = 150). (B) Correlation analysis between the PMN-MDSC signature (Condamine *et al.,* 2016) and the expression of *PADI2* and *PADI4* in the Prostate Cancer Transcriptome Atlas (PCTA), using the PCTA web tool.
